# HIV integrase compacts viral DNA into biphasic condensates

**DOI:** 10.1101/2024.03.15.585256

**Authors:** Pauline J. Kolbeck, Marjolein de Jager, Margherita Gallano, Tine Brouns, Ben Bekaert, Wout Frederickx, Sebastian F. Konrad, Siska Van Belle, Frauke Christ, Steven De Feyter, Zeger Debyser, Laura Filion, Jan Lipfert, Willem Vanderlinden

## Abstract

The human immunodeficiency virus (HIV) infects non-dividing cells and its genome must be compacted to enter the cell nucleus. Here, we show that the viral enzyme integrase (IN) compacts HIV DNA mimetics *in vitro*. Under physiological conditions, IN-compacted genomes are consistent in size with those found for pre-integration complexes in infected cells. Compaction occurs in two stages: first IN tetramers bridge DNA strands and assemble into “rosette” structures that consist of a nucleo-protein core and extruding bare DNA. In a second stage, the extruding DNA loops condense onto the rosette core to form a disordered and viscoelastic outer layer. Notably, the core complex is susceptible towards IN inhibitors, whereas the diffuse outer layer is not. Together, our data suggest that IN has a structural role in viral DNA compaction and raise the possibility to develop inhibitors that target IN-DNA interactions in disordered condensates.

**Teaser:** Single-molecule studies demonstrate the mechanism, dynamics, and drug-susceptibility of viral genome compaction by HIV integrase.

## Introduction

To establish infection, the human immunodeficiency virus (HIV) traverses multiple cellular micro-environments of human host cells, using only a limited arsenal of viral proteins. HIV infects non-dividing cells, and viral particles must enter the cell nucleus to perform integration, thereby irreversibly establishing a provirus in the human genome. While the molecular structures and mechanisms of reverse transcription (*1*,*2*) and integration (*3–7*) are reasonably well understood, the composition and structural assembly of the large nucleoprotein complexes wherein they occur, *i.e.* the reverse transcription complex and the pre-integration complex (PIC) respectively, remain obscure.

Recent work has dramatically impacted our view on the intracellular nature of the virus particle: in contrast to previous belief, HIV transport towards the nucleus occurs within the confines of a (largely) intact viral capsid (*8*, *9*). The new insights suggest that reverse transcription takes place in the confines of the capsid, which in turn requires constraining the increasing mechanical stresses due to the conversion of single-stranded HIV RNA into a much stiffer double-stranded copy-DNA (cDNA) of the HIV genome. Together, the observations suggest the need for a molecular mechanism to compact the DNA within the capsid.

In this context, it was found that the estimated copy-number of the viral protein integrase (IN) in PICs (*10*) is ∼150-250, at least one order of magnitude larger than the number of IN (4-16 copies; (*4*, *11*)) required for intasome assembly and catalysis of integration. This raises the question of whether and how such a large excess of IN is beneficial during the replication cycle. While IN serves a functional role during the later stages of HIV replication, in particular during viral assembly and maturation through binding to viral RNA (*12*), IN mutants defective for non-specific DNA binding are impaired for nuclear import (*13*) and reverse transcription (*14*), suggesting a critical role of IN within the cytoplasmic viral capsid also during the early stages of infection.

Here, we provide evidence for an additional functional role of IN by showing that IN can effectively compact viral DNA mimetics under physiologically relevant conditions. Dimensions of the resulting condensates agree favorably with those of PICs purified from (*15*) or within the infected cell (*16*). Our data, obtained by single molecule force microscopy and spectroscopy techniques, and complemented with coarse-grained polymer simulations, suggest a bridging-induced attraction mechanism (*17–19*) that features an on-pathway “rosette” intermediate preceding full collapse. The fully collapsed condensates retain the mechanically robust rosette core and a diffuse viscoelastic outer layer, reminiscent of current models of eukaryotic genome folding (*20*) – albeit at a length scale orders of magnitude shorter. Importantly, we demonstrate a differential susceptibility towards allosteric IN inhibitors for the rosette folding intermediate *versus* the fully collapsed condensate.

## Results

### IN binds viral DNA ends with moderate selectivity

Binding of IN to viral DNA ends is a prerequisite for the catalysis of the integration reaction. Here, we use atomic force microscopy (AFM) imaging to quantify the selectivity of recombinant IN for the specific ends in short viral cDNA mimics (Fig. 1a), as a function of the level of 3’ processing. From AFM images, we determine the position of the bound IN complexes with respect to the nearest DNA end in IN-DNA nucleoprotein complexes (Fig. 1 and Supplementary Fig. S1), which enables us to estimate the selectivity of end-binding (*21*) 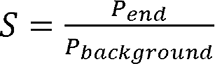. We first investigated binding of IN to a 491 bp cDNA mimic with blunt ends in buffer containing 1 mM Mg^2+^. The distribution of binding events along the internal DNA is essentially homogenous (Supplementary Fig. S1), implying the absence of high affinity internal binding sites. Assuming (*22*) a footprint of 16 bp, there are 476 potential binding sites in the cDNA mimic, and we estimate a selectivity *S* = 64 ± 13 (from *N*=159 molecules; errors were estimated from counting statistics using standard error propagation; Fig. 1g). Next, we studied binding in buffer wherein Mg^2+^ is replaced by Ca^2+^ to impede catalysis of 3’ processing (*23*) and we find a reduced selectivity *S* = 19 ± 7 (*N* = 122; Fig. 1c). Last, we quantify binding to HIV DNA mimetics with 3’ pre-processed ends (see Methods), which results in an increased selectivity, *S* = 256 ± 15 (*N* = 129). Thus, the selectivity of binding to the specific blunt HIV ends increases by more than 10-fold by catalysis of 3’ processing. Nevertheless, binding selectivity overall is only moderate; the probability of binding to a random position in genomic length viral DNA (∼8-9 kbp) is still 1-2 orders of magnitude larger than binding to the specific ends.

**Figure 1.**
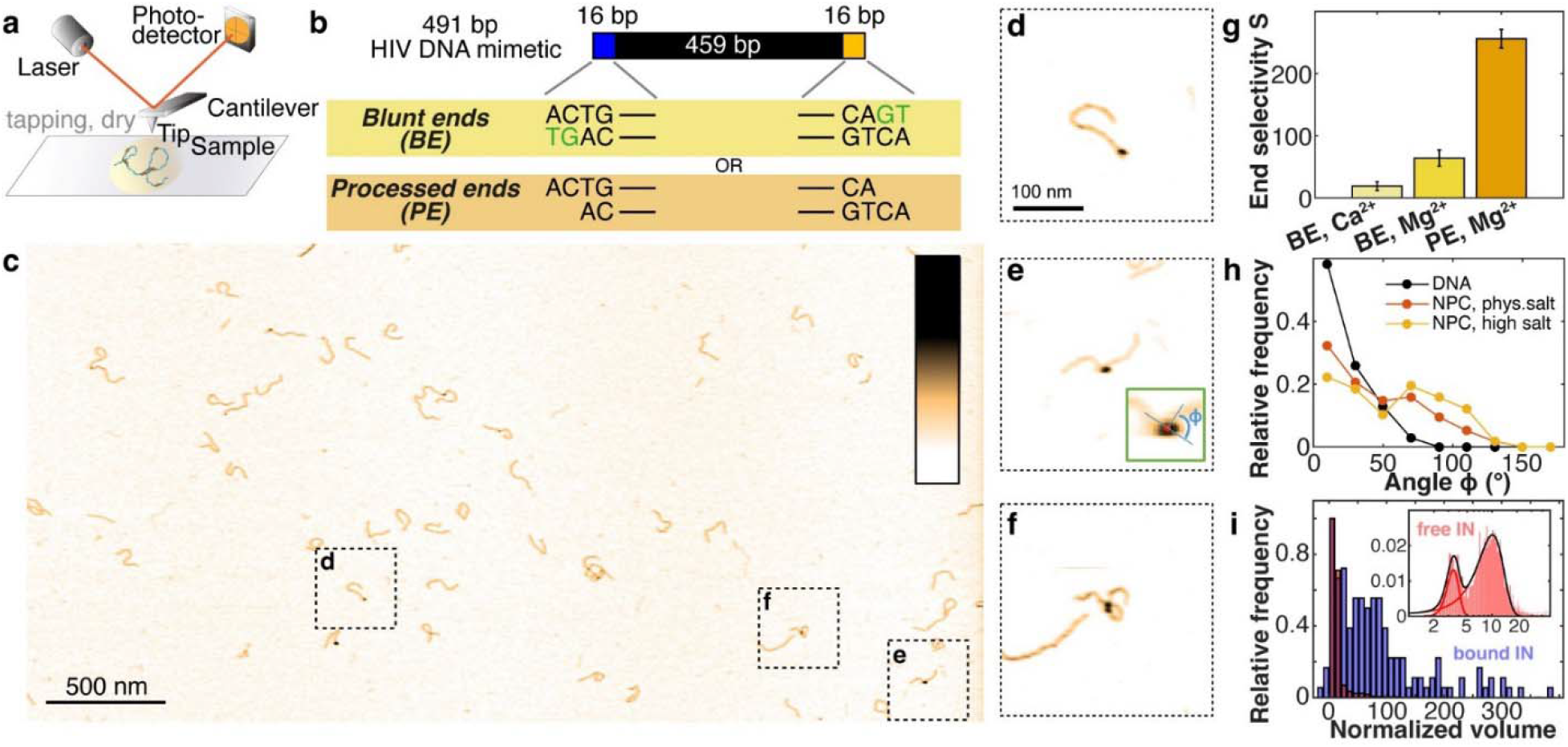
IN specific and aspecific binding to HIV DNA mimetics. **a.** Schematic representation of the experimental set up for AFM imaging experiments. **b.** Schematic of the 491 bp DNA used to mimic the HIV cDNA, both as a blunt ended (BE) or as a 3’-preprocessed substrate (PE). **c.** Overview AFM topograph of the 491 bp DNA in the presence of 100 nM IN. Color range is 1.5 nm. **d.** HIV DNA mimetic with end-bound IN. **e.** HIV DNA mimetic with integrase bound at internal sites and definition of bend angle (inset). **f.** HIV DNA mimetic with integrase-mediated DNA looping. **g**. Selectivity for end-binding over binding to internal sites in the mimetic as a function of 3’ processing and buffer conditions. From left to right: blunt DNA ends, in the presence of Ca^2+^; blunt DNA ends, in the presence of Mg^2+^; processed DNA ends, in the presence of Mg^2+^. The IN binding site size is assumed to be 16 bp. Error bars are based on counting statistics. **h.** Bend angle distribution measured at 10 nm length scale for bare DNA and IN-DNA nucleoprotein complexes (NPCs) deposited from near-physiological salt buffer (50 mM NaCl, 1 mM MgCl_2_, 10 mM Tris-HCl pH = 7.4; *N* = 232) or high salt buffer (250 mM NaCl, 5 mM MgCl_2_, 10 mM Tris-HCl pH = 7.4; *N* = 109). **i.** Histogram of the normalized volume of IN free in solution (red) and IN bound to DNA (blue). In free solution, we find a small fraction of IN monomers and a dominant fraction of IN dimers (inset). For IN bound to DNA, we find the resulting protein volume distribution to be significantly broader and shifted to larger volumes as compared to the DNA-unbound IN population.

### Unspecific IN binding infers DNA bending and looping

As most IN-DNA complexes bind to internal sequences, *i.e.*, non-sequence specifically, we next investigate whether non-specific binding alters DNA structure. We quantify IN-induced DNA bending (Fig. 1e,h) and find that the bend angle distribution is much broader and shifted to larger bend angles at positions where IN is bound, as compared to the distribution of bare DNA bends. At higher ionic strength (250 mM NaCl, 5 mM MgCl_2_), the fraction of complexes with large bend angles (> 50° bending; Fig. 1h) increases with respect to complexes with smaller bend angles, suggesting the presence of (at least) two binding modes: a first mode that is salt-sensitive and minimally invasive, and a second binding mode that is less sensitive to ionic strength and infers strong DNA bending.

Beyond bending, a small fraction (∼5% of all internally bound complexes) forms synapses, i.e., intramolecular DNA loops mediated by IN (Fig. 1f). Using a longer viral DNA mimetic (1000 bp) and otherwise identical conditions (100 nM IN), the fraction of looped DNA increases to 30 ± 5% of all internally bound nucleoprotein complexes (*N* = 93). This increase is consistent with the expectation that looping is facilitated by the longer DNA length, since ∼500 bp corresponds to only ∼4 times the bending persistence length of DNA.

AFM volumetric analysis suggests that IN in the absence of DNA is monomeric and mostly dimeric, whereas in the DNA-bound state it takes on higher oligomers, in particular tetramers (Fig. 1i and Supplementary Fig. S2). Together, our results demonstrate that IN binding to random positions along the DNA is prevalent, induces higher order IN oligomerization, and can alter DNA conformation by bending and looping.

### Two-stage compaction of genomic length viral DNA

To investigate how IN binding affects DNA of lengths similar to the HIV genome, we titrated linearized mini-HIV DNA(*7*) of different lengths (3.4 kbp, 4.8 kbp, 9.1 kbp) with IN and quantify the resulting complexes using AFM (Fig. 2a and Supplementary Fig. S3 to S5). To directly relate the conformations of adsorbed complexes to their solution state, we performed AFM imaging under kinetic trapping conditions, where the resulting images are close to a 3D to 2D projection (*24–26*).

**Figure 2.**
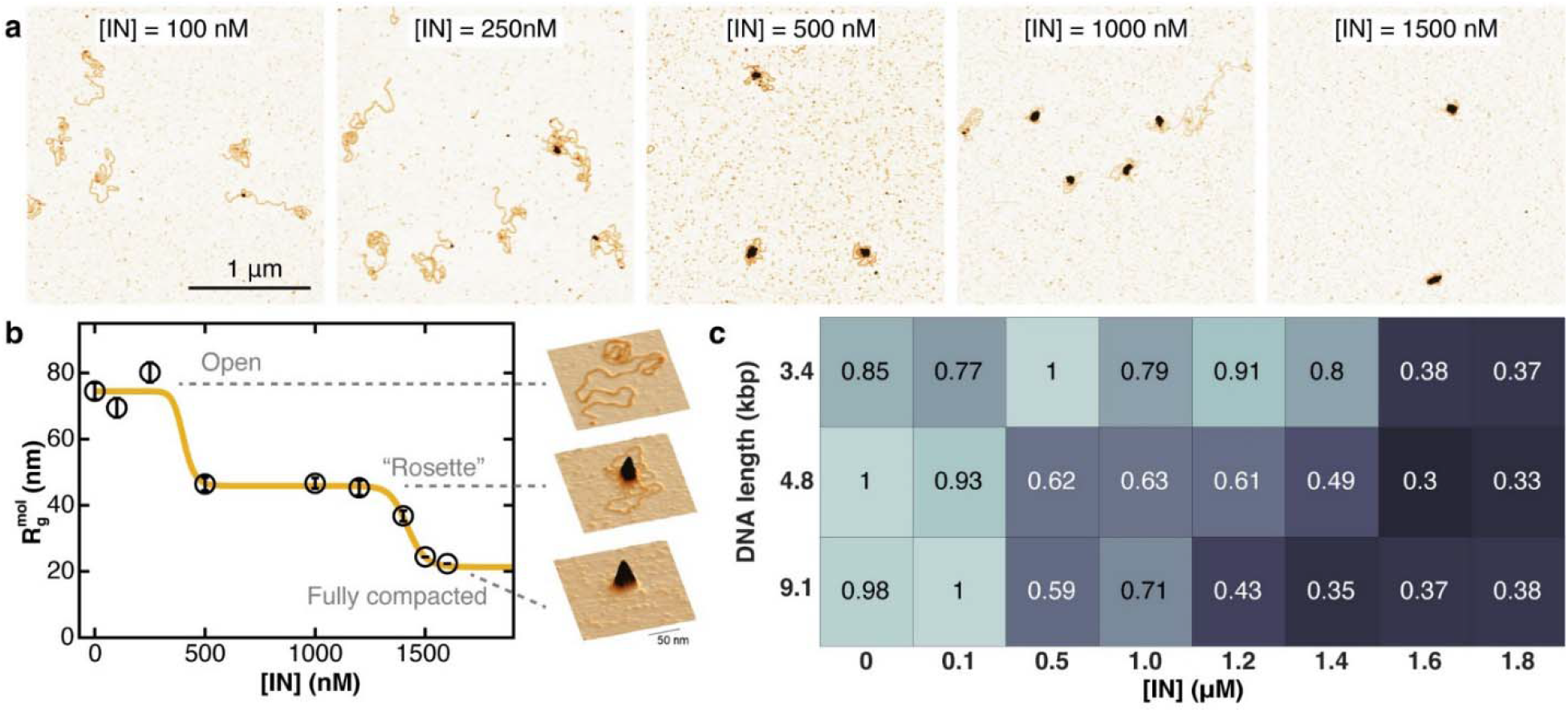
IN compacts long viral DNA mimetics into biphasic condensates. **a.** AFM topographs of 4.8 kbp DNA in the presence of (from left to right) increasing IN concentrations. **b.** Radius of gyration of the nucleoprotein complex for 4.8 kbp DNA at 60 mM ionic strength as a function of [IN]. The data reveal two distinct compaction transition and are fitted by a double-Hill-fit: the first transition from open state to rosette state occurs at ∼ 500 nM, the second transition from rosette state to fully compacted state at ∼ 1100 nM. Representative AFM images are shown as insets on the right. Symbols and error bars are the mean and s.e.m. from typically ≥ 100 molecules per condition. **c.** Phase diagram depicting the normalized radius of gyration of the nucleoprotein complexes formed as a function of [IN] and DNA length, normalized to the largest *R*_g_ observed for each DNA length. With increasing DNA length, the compaction transitions shift to lower IN concentration. Intriguingly, at DNA length > 3.4 kbp a biphasic compaction transition is observed, whereas at 3.4 kbp the transition occurs in one step.

We first used a 4.8 kbp mini-HIV DNA(*7*) for titration with IN in the concentration range [IN] = 0-2 μM. At IN concentrations of < 500 nM, we observe localized IN binding and IN-induced looping. Increasing the concentration to [IN] = 500 - 1200 nM results in the formation of large nucleoprotein complexes featuring a central nucleoprotein core surrounded by bare DNA loops. Because of their appearance, we refer to these complexes as “rosettes”. At even higher IN concentrations (> 1200 nM), the DNA is further compacted, and the nucleoprotein complexes are predominantly in globular conformations wherein individual DNA strands are no longer visible. On occasion, these intramolecular complexes self-associate to form large conglomerates comprising multiple DNA copies (Supplementary Fig. S4f, S5d,e,f).

We quantified DNA compaction upon titration of mini-HIV DNA with IN by automated analysis of the height-weighted 2D radius of gyration 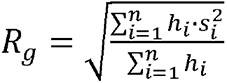 where *s_i_* is the distance of pixel *i* with respect to the molecular center of mass, and *h_i_* is the height value in pixel *i*. Consistent with the qualitative observations of DNA-IN complex geometries, the *R_g_* data reveal that IN significantly compacts DNA in two successive, highly cooperative transitions: a first transition occurs at [IN] ∼ 400 nM and reduces *R_g_* by approximately 40% (from 75 nm at [IN] ≤ 250 nM, to 45 nm for [IN] = 500-1200 nM). A second transition is observed at [IN] ∼ 1400 nM, and further reduces *R_g_* ∼ two-fold (Fig. 2 and Supplementary Fig. S6).

Condensation of nucleic acids with proteins and the formation of nucleoprotein complexes are in general highly dependent on environmental parameters. We mapped out the dependence of IN-induced DNA condensation on environmental conditions and observed that compaction is independent of incubation temperature (we tested 37 °C and room temperature, i.e. 22.5 °C; Supplementary Fig. S7) and incubation time (Supplementary Fig. S8), but strongly dependent on ionic strength(*27*) (Supplementary Fig. S9 and S10). We constructed a phase diagram of condensate formation as a function of integrase and salt concentration (Fig. 2c and Supplementary Fig. S11). The data show that both rosette formation and full collapse are disfavored by high salt concentration but occur under physiologically relevant conditions. As expected from the modest end-binding selectivity (Fig. 1g), we find no effect of the specific miniHIV terminal sequences on compaction efficiency (Supplementary Fig. S12).

Having characterized the compaction behavior of a 4.8 kbp HIV DNA mimic, we next explore how DNA length alters IN-induced condensation. We find that the degree of compaction is independent of the DNA contour length and reduces *R_g_* approximately 3-fold at full compaction compared to bare DNA without IN present (Supplementary Fig. S13). Nevertheless, the critical concentrations for the compaction transitions increases for shorter DNA lengths (Fig. 2c and Supplementary Fig. S6). Most notably, full compaction of the 3.4 kbp HIV mimetic occurs in a single transition (i.e., the rosette state is no longer observed), in contrast to the biphasic behavior observed for 4.8 kbp and 9.1 kbp DNA. The absence of a rosette state with DNA loops for the 3.4 kbp construct is in line with a previously identified DNA length threshold for bridging induced looping (*19*). Last, the radius of gyration of the fully compacted state for 9.1 kbp DNA (approximately the length of the viral genome) is ∼ 32 nm, close to the Stokes’ radius found for purified pre-integration complexes (*15*) (28 nm) and in agreement with the molecular dimensions of pre-integration complexes in infected cells (*16*). Taken together, our quantitative AFM analysis indicates that genomic length viral DNA is compacted via two distinct transitions into condensates with dimensions that are in quantitative agreement with those found in infected cells.

### Simulations suggest bridging-mediated attraction mechanism with a critical role for IN-IN interactions

To obtain a mechanistic understanding of how local interactions lead to DNA compaction, we turn to Monte Carlo (MC) simulations (Fig. 3; Supplementary Methods). We model DNA as a chain of beads, each representing 12 bp (equal to the approximate binding site size of IN) joined by springs. A bending potential between subsequent beads is added to reproduce the experimentally determined bending stiffness of DNA. IN tetramers are similarly modeled as beads of the same size as the DNA beads, with an isotropic binding interaction with the DNA beads. We capture the tetrameric binding nature of the active IN complex by introducing a maximum valence of 4: each IN bead can bind to at most 4 DNA beads (Fig. 3a) (see Supplementary Information). We determine the interaction strength of IN binding to a single DNA segment to be *ε* = 5 *k*_B_*T* (where *k*_B_ is Boltzmann’s constant and *T* the absolute temperature) by direct quantitative comparison (*21*) to experimental data (Fig. 3h). For the IN-IN interaction we consider two cases: (i) no IN-IN interaction (beyond excluded volume interactions) and (ii) IN-IN interactions (i.e. IN-IN attractions in addition to the excluded volume interactions). We perform extensive simulations at different IN concentration for both cases and analyze the DNA-IN conformations as a function of time. During these simulations, the IN concentration is controlled via a grand canonical ensemble (see Supplementary Information) to ensure that the concentration of free IN remains constant. To identify different states of compaction and the pathways between them, we develop a classification pipeline using principal component analysis of 15 different structural parameters and a Gaussian mixture model (Fig. 3b-g and Supplementary Information).

**Figure 3.**
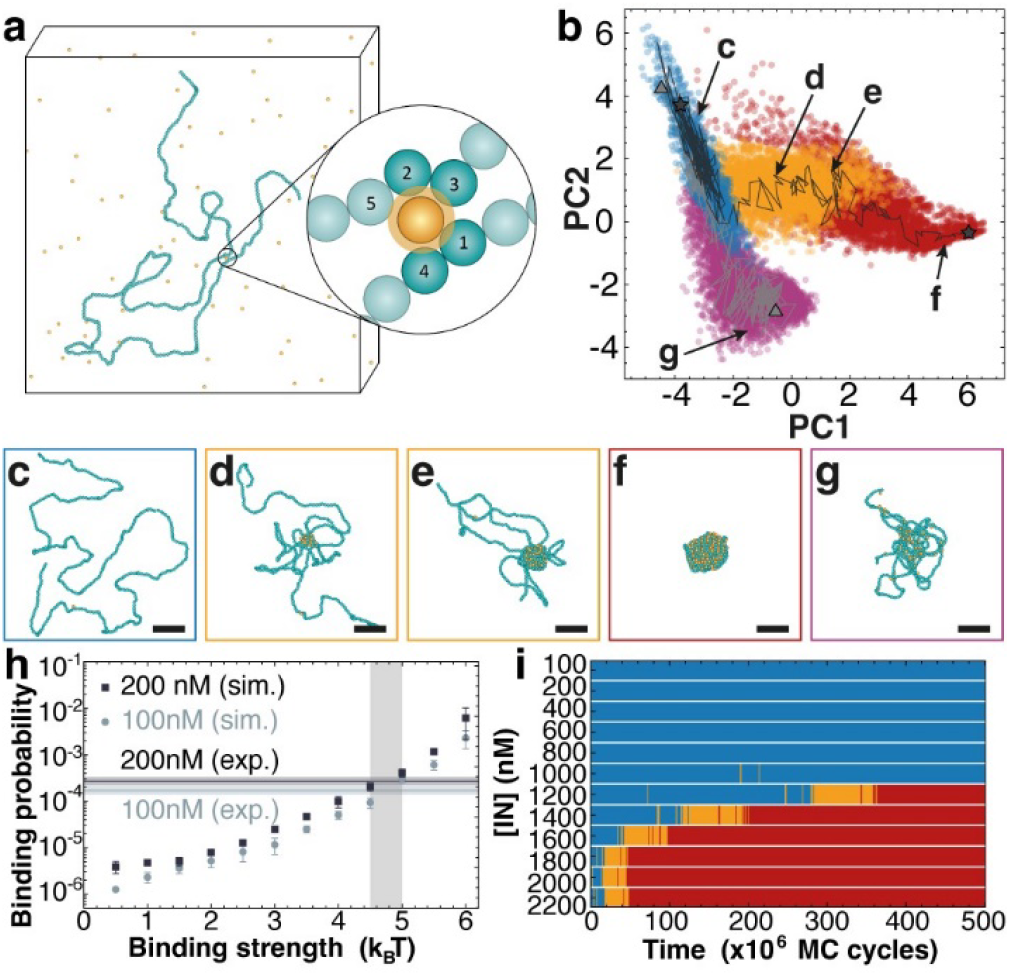
Monte Carlo (MC) simulations show that IN protein-protein interactions are required for biphasic compaction. **a.** Schematic representation of the setup for MC simulations. DNA beads (4 nm) are colored in light blue and IN beads (same size) are colored in yellow. The box size is not to scale. **b.** Principal component analysis separates the large dataset of DNA-IN complexes into several categories of DNA-IN complexes (depicted as different colors, which were assigned using a Gaussian mixture model). The grey trajectories indicate two typical simulation paths of 4.8 kbp DNA (408 beads). Without IN-IN attraction, the left path is taken (triangles at start and end point), here no rosette intermediates are formed. Only if IN-IN attractions are added (right path; stars at start and end point), rosette formation (yellow) as well as full compaction are possible (red). **c-g.** Some typical conformations along the pathways in panel b. The scale bar indicates 40 nm. Free IN are not depicted. **h.** IN-DNA binding probability (mean ± SD from 5 independent simulations) as a function of binding strength (in k_B_T) from MC simulations. The horizontal lines indicate the experimentally determined binding probability for 100 nM and 200 nM IN from AFM images. The crossing area of experiment and simulations allows to estimate the IN-DNA binding energy to be 4.5-5 k_B_T. **i.** Time evolution of the state of compaction (same color as in panel b) of a 4.8 kbp DNA during a MC simulation with IN-IN attractions for varying IN concentrations, well in line with the experimental data (Fig. 2). Each simulation was selected as the most typical one from 12 simulations per IN concentration.

A key result is that we observe distinct pathways in the presence and absence of IN-IN attractions: MC simulations that only include IN-DNA binding (*17*), but no IN-IN attraction, exhibit DNA compaction, but fail to reproduce the two-phase compaction and the experimentally observed rosette conformations. In contrast, if IN-IN attraction is included (*18*) (Fig. 3i), we observe transitions and structures that are in good agreement with the experimental observations (Fig. 2). In summary, our simulations demonstrate that a simple model that includes both IN-DNA and attractive IN tetramer-tetramer interactions can semi-quantitatively reproduce the observed DNA compaction.

### IN-DNA condensates feature a rigid core surrounded by a soft coat

To further characterize the IN-DNA condensates and to probe their mechanical properties that likely play a role in the interior of the capsid and/or during nuclear entry, we use AFM force-volume mapping (*28*, *29*). In this modality, a full force-distance curve (i.e., an approach and retract cycle) is recorded in every pixel of the image (Fig. 4a). Localized interaction forces on the tip are recorded as it moves towards and away from the sample surface (Fig. 4b) and can be directly correlated to the sample topography. We analyze the approach force curves to extract the indentation stiffness *k*, the retract curves to quantify elastic deformation energy *E_Elast._*, and the hysteresis between both curves to obtain a measure of plastic deformation *E_Plast._* (Fig. 4a, inset and Supplementary Fig. S14). From the elastic and plastic contributions to the deformation energy, we can quantify the dimensionless viscoelasticity index(*30*) 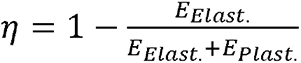, where *η* = 0 for a perfectly elastic deformation and *η* = 1 for a completely viscous deformation, i.e. where the bonds broken during indentation are not reformed during the stress release.

**Figure 4.**
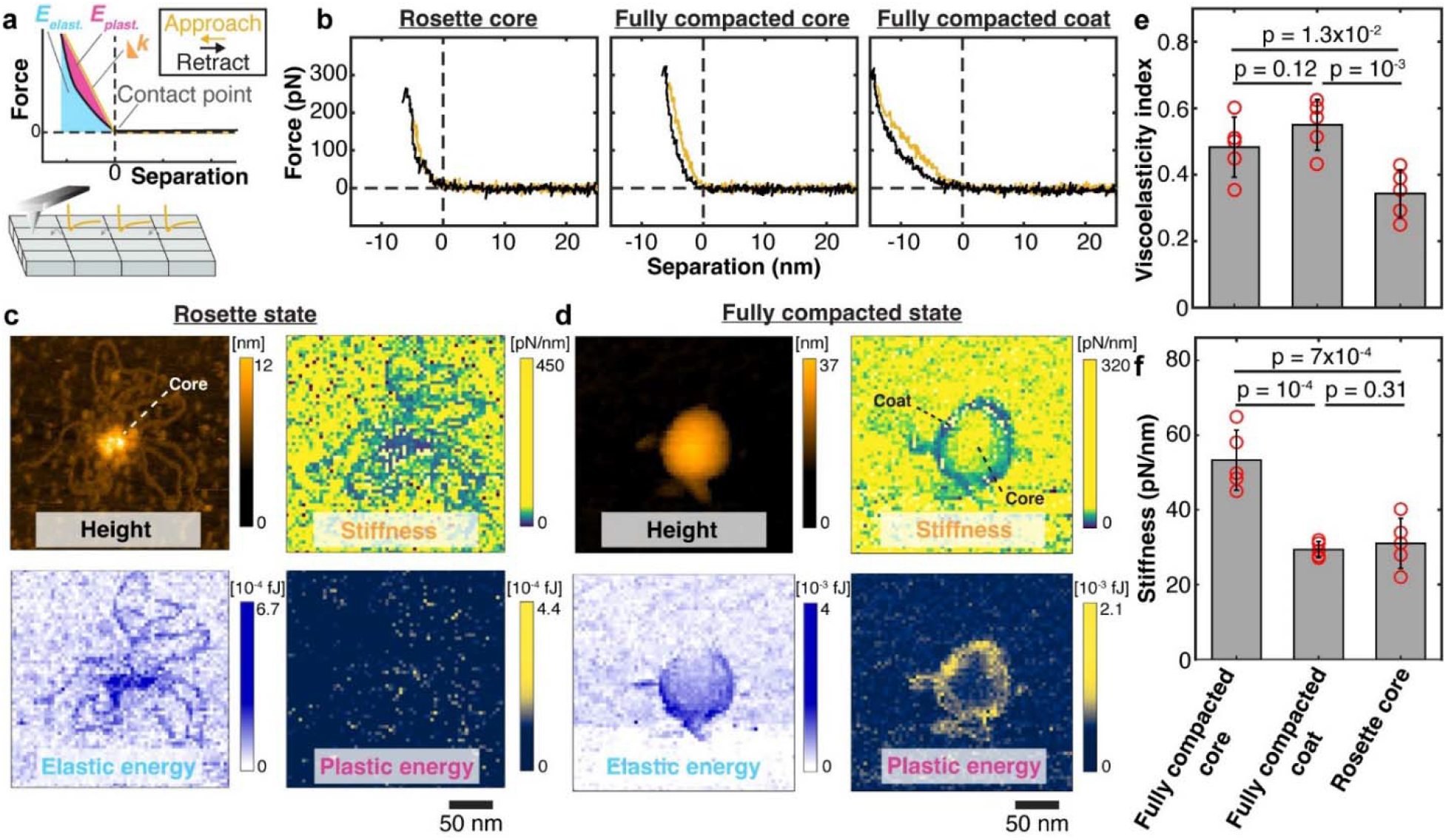
AFM force-volume based multiparametric imaging of IN-DNA condensates. **a.** Schematic depiction of AFM-based force-volume mapping, and derived parameters *k* (indentation stiffness), *E*_plast._ (plastic deformation energy), and *E*_elast._ (elastic deformation energy). **b.** Example force distance curves (yellow: approach; black: retract). **c.** Multiparametric imaging of a condensate in the rosette state: zero-force height (top left), stiffness (top right), elastic energy (bottom left), plastic energy (bottom right). **d.** Multiparametric imaging of a DNA-IN condensate in the fully compacted state, indicating a soft “coat” surrounding a rigid “core”. **e.** Viscoelasticity index (mean ± SD) of the core and coat of fully compacted state, and of core of rosette state. *p*-values are based on 2-sample Kolmogorov-Smirnov test. **f.** Indentation stiffness (mean ± SD) of the core and coat of fully compacted state, and of core of rosette state. *p*-values are based on 2-sample Kolmogorov-Smirnov test.

We first perform force-volume mapping on complexes in the rosette state (Fig. 4c). The approach and retract curves are uniform across the nucleoprotein core and exhibit little hysteresis. The fitted values of *k*, *E_Elast._*, and *E_Plast._* from at least 100 approach and retract curves are pooled and averaged per complex. Averaging over different rosette complexes, we find a mean stiffness *<k>* = 31 ± 7 pN/nm (mean ± SD from *N* = 5 complexes) and a mean viscoelasticity index *<η>=* 0.34 ± 0.07 (Fig. 4e,f).

In a second set of experiments, we use force-volume mapping on fully collapsed condensates (Fig. 4d; *N* = 5). In contrast to force curves recorded on rosettes, we consistently observe substantial hysteresis between the approach and retract curves. Furthermore, we find the overall slopes of the force curves to vary depending on their radial position: at constant peak forces, larger indentations are observed near the edges of the condensates. Indentation curves do not show breakthrough events, and because one would expect decreased indentations due to finite AFM tip size effects near steep topographic features we interpret the spatially resolved features in the condensates to result from complex material properties.(*31*, *32*) The spatially resolved stiffness map is used to identify a “core” and outer layer or “coat”. We pool force curves in the core and find *<k>* = (53 ± 8) pN/nm (error is SD) and a mean viscoelasticity index *<η>=* 0.48 ± 0.09 over the different condensates (Fig. 4e,f). An analogous treatment for force curves recorded on the condensate coats gives *<k>* = (29 ± 2) pN/nm (error is SD), significantly less than the stiffness of the rosette, and a mean viscoelasticity index *<η>=* 0.55 ± 0.08.

We conclude that the fully collapsed condensates feature a relatively rigid, stiff core and more viscous, softer outer layer. The observed stiffness of the fully compacted condensates are on the same order of magnitude, but still significantly lower (by at least ∼5-fold) than the reported stiffnesses of full virus particles (*33*), consistent with the possibility that they could remain in the capsid during transport to and into the nucleus. The pronounced viscoelasticity and hysteresis between approach and retract curves suggests that intrinsic dynamics and re-arrangement of bonds in the complex are much slower than the time scale of the force application and release by AFM (∼5 ms).

### Force spectroscopy of IN-DNA reveals heterogeneous dynamics of compacted loops

To investigate the dynamics and stability of IN-DNA condensates, we performed magnetic tweezers (MT) force spectroscopy experiments. In our assay, we tethered long linear DNA molecules (21 kbp) between the flow cell surface and superparamagnetic beads (Fig. 5a). Using external magnets, we can apply precisely calibrated stretching forces to the tethered molecules (*34*). To investigate the forces involved in IN-DNA condensation, we first introduced IN (2 μM) in the flow cell to interact with tethered DNA molecules under very low stretching forces (F < 0.01 pN) for 10 min, to allow for condensate formation, and then increase the applied forces to mechanically probe the stability and structure of the condensate (Fig 5b). In a first set of experiments, we gradually increase the force from 0.1 to 5 pN with plateaus of constant force of 60 min (0.1 to 0.9 pN) or 30 min (1 to 5 pN), respectively (Fig 5b and Supplementary Fig. S15), while tracking the extension of the DNA tether. In the presence of IN, the DNA is shortened, compared to control experiments in the absence of IN and to the prediction of the WLC model (red line in Fig 5b), in particular in the force range 0.5 – 1 pN (Fig. 5b, inset). At very low forces (< 0.5 pN), we observe slow increases and decreases in DNA extension at constant force (Fig. 5c and Supplementary Fig. S15c-f), suggesting that the breakage and re-formation of interactions is in equilibrium under these conditions, pointing towards a dynamic and reversible nature of the interactions involved in loop compaction. Notably, the largest deviations from bare DNA behavior are observed in the same force range in force-extension experiments (Fig 5b, inset). The observed reversibility of compaction in the MT is consistent with the observation that rosette formation and compaction are reversible in AFM imaging experiments upon dilution.

**Figure 5.**
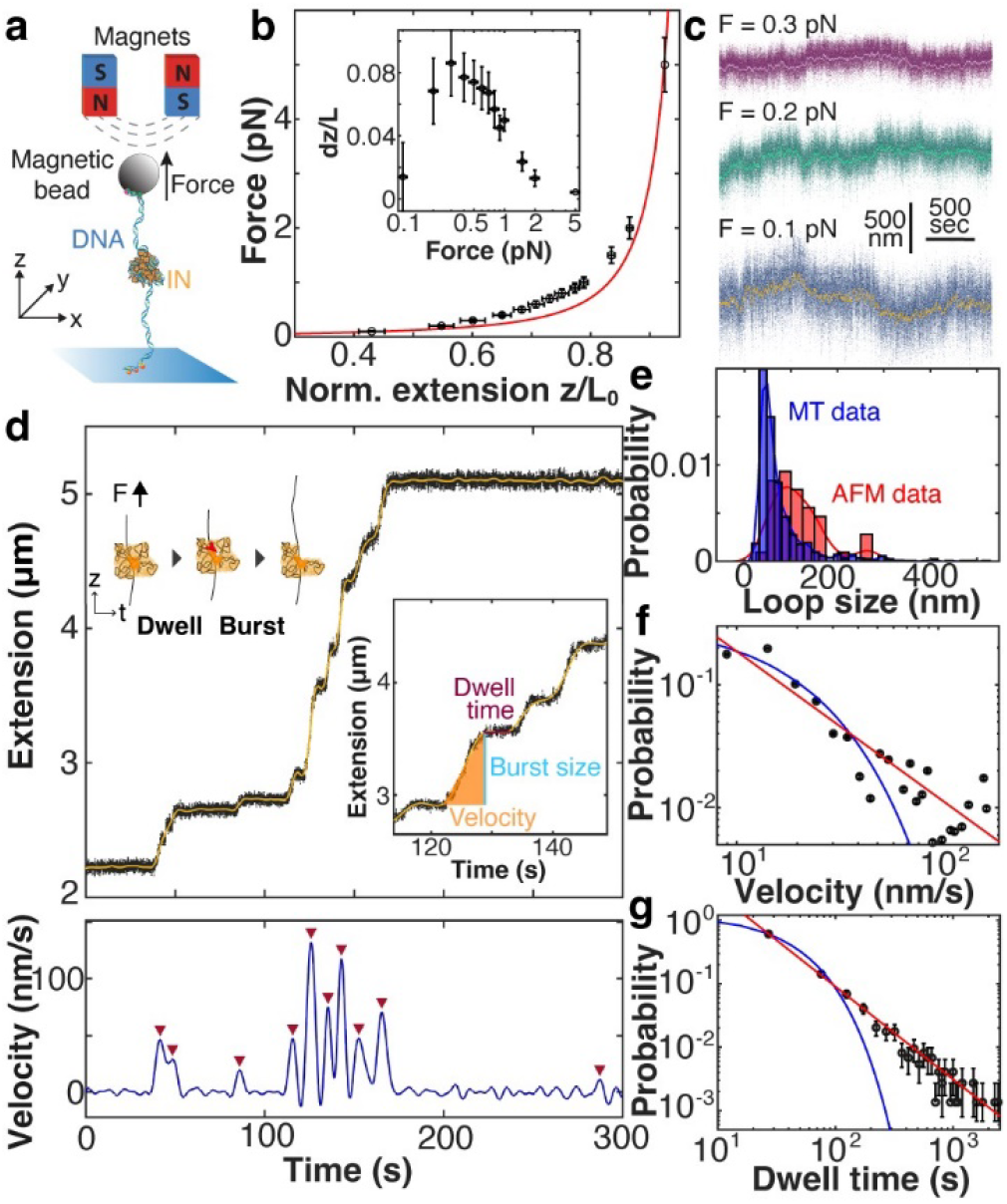
Single-molecule force spectroscopy quantifies dynamics condensate unfolding. **a.** Schematic depiction of the MT force spectroscopy measurements. 21 kbp DNA is tethered between a flow cell surface and magnetic beads and (partially) compacted by IN. External magnets enable the application of precisely calibrated stretching forces. **b.** Force-extension relation at forces ≤5 pN. Extension *z* is normalized to *L_0_*, the extension of bare DNA at *F* = 5 pN. Red line is a worm-like chain (WLC) fit of bare DNA. The inset shows the deviation of the experimental data (mean ± SD) in the presence of IN from bare DNA WLC behavior; the maximum deviation occurs at ∼ 0.3 pN. **c.** Extension-time traces of IN-DNA condensates at constant forces ≤0.3 pN exhibiting dynamic compaction and extension. **d.** Top: Extension-time trace following a force jump from 0.01 to 1 pN. Raw data (68 Hz) are depicted in black, yellow line is smoothed data (Butterworth filter, see Methods and SI for filter parameters). Insets depict a schematic of the events governing dwells (corresponding to extension plateaus), and extension bursts following dissociation of IN-DNA bridge in the core of the complex (red arrow). Bottom: Velocity-time trace, as obtained by differentiation of the filtered extension trace. Red marks indicate extension bursts detected by thresholding. **e.** Burst size distribution (blue bars; *N*= 777 events from 41 independent DNA tethers). For comparison, the loop size distribution obtained by AFM imaging are shown (red bars). A two sample *t*-test suggest no significant difference between both distributions (2-sample t-test; *p* = 0.5) **f.** Weighted bursts velocity distribution fitted with exponential decay (blue line; fitted decay constant 12 ± 3 nm/s) and power law (red line; scaling exponent –1.9 ± 0.3) for 1 nm/s < *v* < 100 nm/s. **g.** Distribution of dwell times between extension bursts and fits using an exponential decay (blue line; decay time of 24 ± 3 s) and power law (red line; scaling exponent –1.0 ± 0.1). Errors reflect 95% confidence intervals on the fit parameter.

In a second set of experiments, we again incubated tethered DNA molecules at a very low force (F < 0.01 pN) in presence of 2 μM IN for 5 min, to allow for the formation of IN-DNA condensates. Then, we apply a force-jump to a constant stretching force of 1 pN. We observe that the extension only increases very gradually – in contrast to control experiments with bare DNA, where the DNA extension increases essentially instantaneously (≤1 s) to the full extension of DNA at the given force (Supplementary Fig. S16). During force-induced extension in the presence of 2 μM IN, we observe extension increments or “bursts”, interspersed by long (>min) pauses or “dwells” where extension remains constant (Fig 5d, top panel). We quantify the extension time traces at constant force by determining i) the dwell times between extension events and for each extension event, ii) the total increase in tether length for the event, and iii) the extension velocity (Fig 5d, bottom panel and Supplementary Fig. S17 to S19).

The increase in length per extension event is in reasonable agreement with the loop sizes found in AFM measurements with short DNA (Fig. 1 and Fig. 5e), suggesting that individual extension events correspond to the release of compacted loops. The extension velocities are much smaller (by at least two orders of magnitude) than what would be expected for the release of free DNA loops (*35*, *36*) and show a broad velocity distribution (Fig. 5f), suggesting that IN-compacted DNA loops are released under forces of ∼1 pN, but that the release is slowed down by protein interactions that cause an effective friction and exhibit heterogeneous dynamics, consistent with the viscous nature of the soft coat region observed by AFM probing. The distribution of dwell times between extension events has a mean lifetime of (24 ± 3) s and does not follow a single exponential, but rather a power law with scaling exponent −1.0 ± 0.1 (Fig. 5g), suggesting heterogeneous interactions and dynamic disorder (*37*).

The finding that extension plateaus (characterized by long dwell times ∼1 min) separate bursts of extension increments (which occur on the ∼s timescale), suggests that force is first exerted on mechanically stable and long-lived DNA-protein bonds that shield the hidden length in compacted loops from opening up. Once these long-lived bonds break, compacted loops are released and open under the applied force, but still experience friction originating from protein-protein and protein-DNA interactions. Together, our MT data further confirm the biphasic nature of fully compacted condensates, and indicate heterogeneous interactions with high levels of dynamic disorder.

### Allosteric IN inhibitors interact differentially with rosettes versus collapsed condensates

Our experimental data suggest that interactions in the core are mediated by IN tetramers and higher order assemblies thereof (Fig. 1i). As IN tetramers are the preferred target for allosteric integrase inhibitors (*38*), we tested the susceptibility of condensates in the rosette and fully compacted states towards the small molecule, quinoline-derivative inhibitor CX014442, an integrase inhibitor originally developed to inhibit association with the coactivator LEDGF/p75 (LEDGIN; this class of inhibitors has also been referred to as ALLINI for *allosteric integrase inhibitor*) (*39*). In the following, we will refer to CX014442 from now on as the “inhibitor” for simplicity.

In a first set of experiments, we introduced the inhibitor at 1 and 2 μM, at fixed [IN] = 1 μM, i.e. under condition where the rosette is formed, and studied the resulting complexes via AFM imaging (Supplementary Fig. S20). At equimolar [inhibitor]:[IN] ratio, the typical single nucleoprotein clusters of the rosette conformations appear partly disintegrated (Fig. 6a). We observe an increase in the radius of gyration of the entire molecule, albeit statistically not significant (*p* = 0.35; Fig. 6b). In contrast, the radius of gyration of the core decreases significantly (*p* = 0.024; Fig. 6c and Supplementary Fig. S21). At 2:1 [inhibitor]:[IN] ratio, the rosette core is disintegrated completely: individual IN complexes are still bound along the DNA contour but do not exhibit bridging interactions. Consistently, we find significant changes in cluster size and integrity (via *R_g,core_*; *p* = 10^-25^), as well as significant changes in compaction of the condensate (via *R_g,mol_*; *p* = 2·10^-14^).

**Figure 6.**
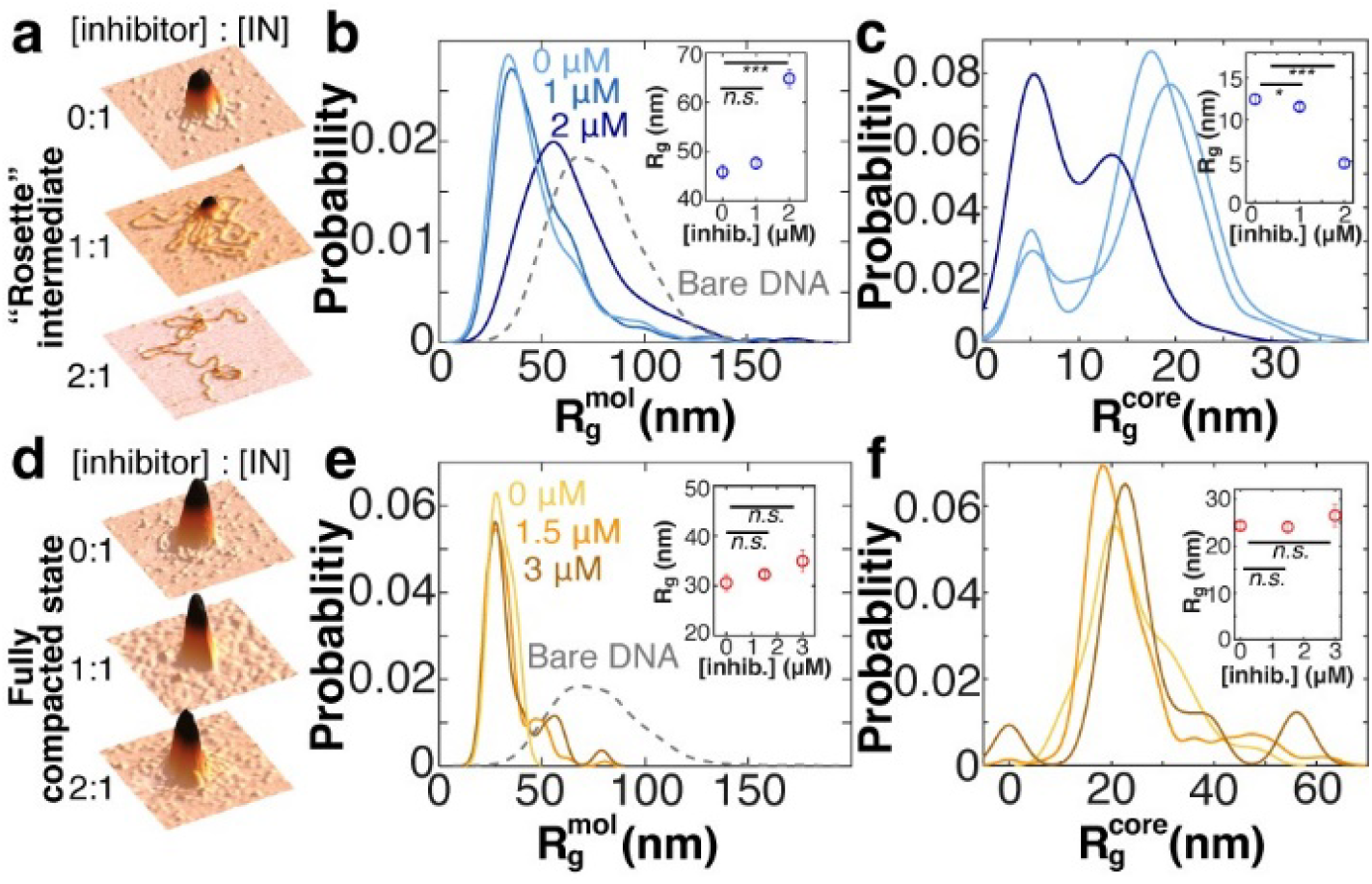
Allosteric inhibitors affect compaction behavior of long viral DNA mimetics. **a.** AFM topographs of the 4.8 kbp DNA in the presence of 1000 nM IN (rosette state) and no, 1000 nM, 2000 nM inhibitor. The inhibitor was added after complex formation. **b.** *R_g_*-distributions for the entire molecule at 1000 nM IN and no, 1000 nM, 2000 nM inhibitors. As a reference, the distribution for bare DNA is shown as well. With increasing inhibitor concentration, less compaction is observed. **c.** Same conditions and color coding as in panel b only the *R_g_*-distributions for the core are shown. **d.** AFM topographs of the 4.8 kbp DNA in the presence of 1500 nM IN (fully compacted state) and no, 1500 nM, 3000 nM inhibitor. The inhibitor was added after complex formation. **e.** *R_g_*-distributions at 1500 nM IN and no, 1500 nM, 3000 nM inhibitors. As a reference, the distribution for bare DNA is shown as well. **f.** Same conditions and color coding as in panel e only the *R_g_*-distributions for the core are shown. Even at high inhibitor concentrations, the fully compacted DNA-IN complexes do not disassemble under influence of the inhibitor. The insets in panels b, c, e, and f shows the means ± SEM of the respective distributions. Significance values are based on 2-sample Kolmogorov-Smirnov test: * indicates *p <* 0.05; *** indicate *p* < 0.001, *n.s.* indicates not significant.

In a second set of experiments, we introduced the inhibitor at 1.5 and 3 μM, at a fixed [IN] = 1.5 μM, i.e. to fully compacted condensates (Fig. 6d). Strikingly, independent of the inhibitor concentration, the nucleoprotein assemblies retain their fully compacted conformations akin to those found in the absence of inhibitor and we observe no significant changes in cluster size (Fig. 6e,f). We note that the influence of the inhibitor on rosette or fully compacted conformations does not depend on order of addition, indicating that steric effects on inhibitor binding cannot explain our observations (Supplementary Fig. S21).

In general, LEDGINs have multiple mechanisms of action during HIV replication. By interference with the binding of LEDGF/p75 they redirect integration into less optimal sites. LEDGINs also allosterically interfere with the enzymatic activity of integrase and induce aberrant multimerization (*40*). Aberrant multimerization of IN during virus assembly results in defective virus particles. LEDGINs do affect IN-IN interactions and, in particular induce the loss of IN into inactive aggregates at high IN concentrations (*12*, *41*). Consistent with this view, we observe protein aggregates by AFM imaging (Supplementary Fig. S20). However, the fact that at 1.5 μM IN no visible changes on fully compacted nucleoprotein complexes are observed in the presence of inhibitor strongly suggests that the major effect on IN-induced condensation is not due to the loss of IN in inactive aggregates. In addition, the differential effects of the inhibitor on rosette formation and the fully collapsed state indicate that the protein-protein and/or protein-DNA interactions involved are different for the different compaction stages.

## Discussion

Here we use single molecule experiments in combination with Monte Carlo simulations to analyze nucleoprotein condensates formed between viral cDNA mimics and recombinantly expressed HIV IN. IN induces (flexible) DNA bends and mediates the formation of DNA loops. Furthermore, DNA-bound IN complexes were found to be larger than unbound complexes, suggesting that DNA binding induces conformational changes leading to higher order oligomerization (Fig. 7). At concentrations 500-1000 nM, IN induces formation of large DNA complexes with multiple DNA loops extending from the center that we refer to as rosettes. At higher IN concentrations 1000-1500 nM, a second compaction transition occurs in which the extruding loops of the rosette structure collapse onto the core of the condensate and full compaction is achieved. The resulting fully collapsed condensates drastically (∼30-fold) reduce the volume occupied by the viral genome. In addition, our data indicate that in the fully collapsed condensate, a rigid central core is surrounded by a diffuse outer layer with relaxation times on the ∼min timescale. It is interesting to note that this architecture resembles a polymer brush, the model that similarly describes the folding of eukaryotic chromosomes (*42*) that are orders of magnitude larger and also plays key roles in the organization of prokaryotic genomes (*43*). Importantly, we experimentally demonstrate that the center of the rosette and the outer layer of the fully compacted IN-DNA complexes interact differently with the inhibitor CX014442: while the center of the rosette dissolves in an inhibitor concentration-dependent manner, the outer layer of the fully compacted state is not susceptible to the inhibitor in the concentration range probed, irrespective of the order of addition of the inhibitor (Supplementary Fig. S21). Intriguingly, this drug-susceptibility (Fig. 6) is in stark contrast to the mechanical (Fig. 4 and 5) and thermodynamic stability (Fig. 2).

**Figure 7.**
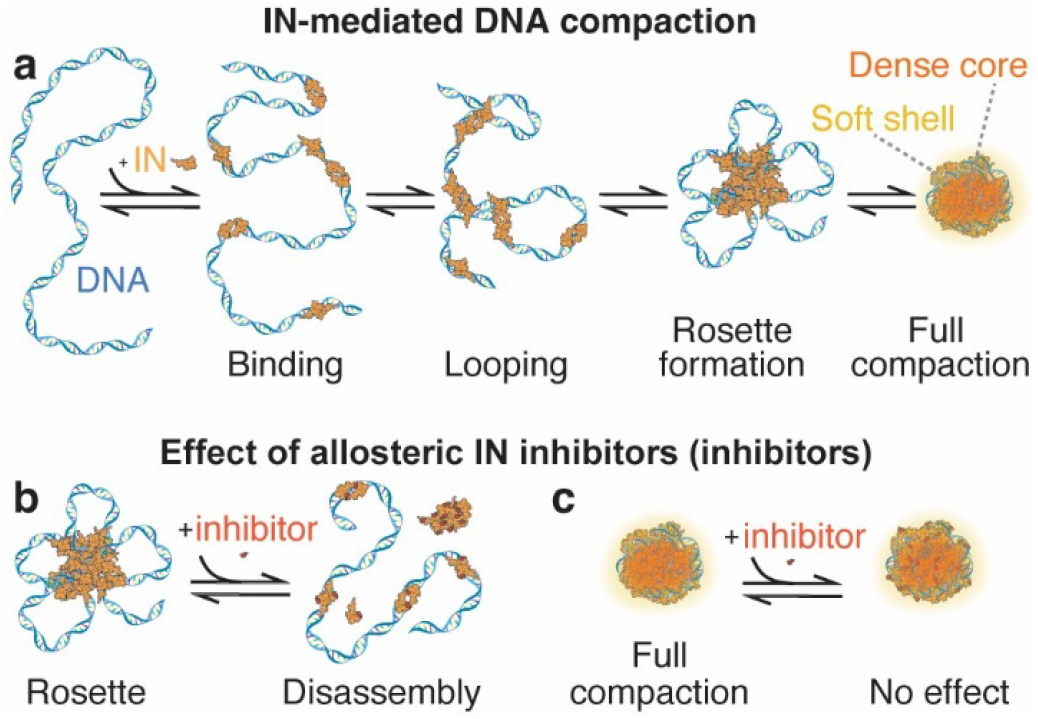
Model for IN-mediated DNA compaction. **a.** Schematic model of DNA compaction by IN. At increasing IN concentrations, the protein binds to the DNA and can form DNA loops. At intermediate concentration, loop-extruding complexes, so-called rosettes, form. At high IN concentrations, full compaction occurs. **b.** The allosteric IN inhibitor CX014442 can de-compact rosettes. **c.** The allosteric IN inhibitor CX014442 cannot de-compact fully compacted DNA-IN condensates. We observe that all compaction and de-compaction processes are reversible.

Provided that our experiments give a very reductionist view of HIV DNA genome compaction, it is important to evaluate their potential relevance with regards to viral condensates in the infected cell. In this regard, the size of IN-DNA condensates in vitro compare very favorably to sizes found by super-resolution microscopy in vivo and to those of PICs purified from infected cells (*16*). In addition, IN plays a crucial role during capsid uncoating in vivo (*44*) and IN mutants defective for non-specific DNA binding are impaired for nuclear import (*13*) and reverse transcription (*14*). Thus, it appears that IN plays a critical role within the cytoplasmic viral capsid during the early stages of infection. Maybe, these effects involve an architectural role for IN as a DNA compaction agent – similar to the established role of IN during folding of the RNA genome in nascent virus particles (*12*). Nevertheless, we stress that our data do not rule out the involvement of other DNA interacting viral proteins that have been shown to induce DNA compaction in vitro, such as the nucleocapsid (*45*, *46*) and VpR proteins (*47*).

Finally, we are convinced that the highly quantitative approaches presented here will be instrumental in future efforts to unravel the molecular mechanisms governing retroviral nucleoprotein condensation, e.g. to further establish the role of IN and other DNA-binding viral proteins in compaction of viral DNA. We especially anticipate breakthrough insights into PIC architecture by combining bottom-up (i.e. reconstitution) and top-down (i.e. purification) approaches. Similarly, it would be intriguing to use similar methods to probe the effects of IN on HIV RNA compaction, which has an established biological role during virus maturation (*12*). Last, we expect our experimental platform to be valuable for the in-depth mechanistic investigation of strand transfer and allosteric inhibitors of IN on DNA compaction, and potentially uncover the mode of action of future inhibitors targeting disordered phases of IN and its condensates.

## Materials and Methods

### HIV-1 integrase expression and purification

For all experiments, we used recombinantly expressed C-terminal or N-terminal His-tagged HIV-1 integrase as described previously.(*48*) The small-molecule allosteric IN inhibitors CX014442 were obtained from KU Leuven (Cistim-CD3).

### AFM imaging with short DNA

We used two different short DNA constructs: a blunt-end construct, and a 3’ pre-processed construct. For the blunt-end 491 bp viral cDNA mimic, we used a custom-synthesized double-strand DNA (gblock, IDT; see below) without further purification. For the 3’ pre-processed HIV DNA mimetic, a gblock of 750 bp (gblock, IDT; see below) was first PCR amplified with forward primer 5’-ACT GGA AGG GCT AAT TCA CTC −3’ and reverse primer 5’-ACT GCT AGA GAT TTT CCA CAC −3’. 2 µg of this DNA was cut with the restriction enzymes EcoRI and XhoI. This fragment is used in a ligation reaction with an U3 and U5 pre-processed fragment. The U3 pre-processed fragment was generated by first hybridizing the oligo’s: 5’- /5Phos/TC GAG TTT TAG TCA GAG TGA ATT AGC CCT TCC A −3’ and 5’-ACT GGA AGG GCT AAT TCA CTC TGA CTA AAA C −3’ and subsequently cutting it with the restriction enzyme XhoI and the U5 pre-processed fragment was generated by hybridizing the oligo’s: 5’- /5Phos/AA TTC TTT TAG TCA GTG TGG AAA ATC TCT AGC A −3’ and 5’-ACT GCT AGA GAT TTT CCA CAC TGA CTA AAA G −3’ and subsequently cutting it with the restriction enzyme EcoRI. The ligated fragment was run on a gel and the correct band was cut out and eluted from the gel.

The AFM samples are prepared by incubating 2.5 ng/ μL viral cDNA mimics (491 bp with 16 bp terminal sequences corresponding to viral U3 and U5 ends; see sequence below) and recombinant (C-terminal His-tag) HIV integrase (IN) at different concentrations, in high sodium buffer (10 mM Tris-HCl pH = 7.6; 250 mM NaCl; 5 mM MgCl_2_) and at 37° C for 30 min, and subsequent 5-fold dilution in low sodium buffer (10 mM Tris-HCl pH= 7.6) just prior to drop-casting onto poly-L-lysine-modified mica(*49*, *50*). Additional measurements were performed in the absence of Mg^2+^, using a high salt buffer comprising CaCl_2_ (10 mM Tris-HCl pH= 7.6; 250 mM NaCl; 5 mM CaCl_2_). After allowing adsorption for 30 s, mica substrates were rinsed with milliQ water and dried using a gentle flow of filtered N_2_ or Argon, respectively. AFM imaging was performed in amplitude-modulation (“tapping”) mode in air using a Nanoscope IV Multimode AFM.

#### Sequence of the 491 bp viral cDNA mimic

ACTGG AAGGG CTAAT TCACT CGGGC GAATT CGAGC TCGGT ACCCG GGGAT CCTCT AGAGT CCATG CAAGC TTGAG TATTC TATAG TGTCA CCTAA ATAGC TTGGC GTAAT CATGG TCATA GCTGT TTCCT GTGTG AAATT GTTAT CCGCT CACAA TTCCA CACAA CATAC GAGCC GGAAG CATAA AGTGT AAAGC CTGGG GTGCC TAATG AGTGA GCTAA CTCAC ATTAA TTGCG TTGCG CTCAC TGCCC GCTTT CCAGT CGGGA AACCT GTCGT GCCAG CTGCA TTAAT GGAGG CGGTT TGCGT ATTGG GCGCT CTTCC GCTTC CTCGC TCACT GACTC GCTGC GCTCG GTCGT TCGGC TGCGG CGAGC GGTAT CAGCT CACTC AAAGG CGGTA ATACG GTTAT CCACA GAATC AGGGG ATAAC GCAGG AAAGA ACATG TGAGC AAAAG GCCAG CAAAA GGCCA GGAAC GTGTG GAAAA TCTCT AGCAG T

#### Sequence of the 750 bp gblock

ACTGGAAGGGCTAATTCACTCTGACTAAAAGGATCCCAGTATTTGGTATCTGCGCTCTGCTGAAGCCAGTTACCTTCGGAAAAAGAGTTGGTAGCTCTTGATCCGGCAAACAAACCACCGCTGGTAGCGGTGGTTTTTTTGTTTGCAAGCAGCAGATTACGCGCAGAAAAA AAGGATCTCAAGAAGATCCTTTGATCTTTTCTACGGGGTCTGACGCTCAGTGGAACGAAAACTCACGTTAAGGGATTTTGGTCATGAGATTATCAAAAAGGATCTTCTCGAGGATACAGGATTAGCAGAGCGAGGTATGTAGGCGGTGCTAAAGTATATATGAGTAAACTTGGTCTGACAGTTACCAATGCTTAATCAGTGAGGCACCTATCTCAGCGATCTGTCTATTTCGTTCATCCATAGTTGCCTGACTCCCCGTCGTGTAGATAACTACGATACGGGAGGGCTTACCATCTGGCCCCAGTGCTGCAATGATACCGCGAGACCCACGCTCACCGGCTCCAGATTTATCAGCAAAGCTTCAGCCAGCCGGAAGGGCCGAGCGCAGAAGTGGTCCTGCAACTTTATCCGCCTCCATCCAGTCTATTAATTGTTGCCGGGAAGCTAGAGTAAGTATCTAGACAGTTAATAGTTTGCGCAACGTTGTTGCCATTGCTACAGGCATCGTTCGCTCACGCTCGTCGTTTGGTATGGCTTCATTCAGAATTCTTTTAGTCAGTGTGGAAAATCTCT AGCAGT

### AFM volume analysis of proteins

To estimate the molecular weight of protein particles, we constructed a calibration curve of protein volumes as a function of known protein molecular weights. To correct for the finite size of the AFM tip, we normalize the measured volumes by the volume per nanometer length of co-adsorbed DNA. The normalized volumes of protein particles measured by AFM are well-described by a power law fit: 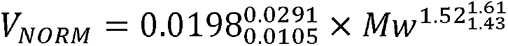 (for *Mw* ≤ 632 kDa; *χ*_Red_^2^ = 1.14; the sub- and superscript numbers indicate the 95% confidence intervals from the fit). To quantify to the volume of DNA-bound proteins, we quantified the volume of the entire protein-DNA complex and subtracted the average volume of surrounding free DNA molecules. Volume normalization using free DNA as a fiducial marker is done as descried above.

### AFM imaging with long DNA

As viral cDNA mimic, we used the miniHIV plasmid, which has been described previously.(*7*) Using the 4751 miniHIV plasmid(*7*) as a parental plasmid, we generated two other viral cDNA mimics (3437 bp and 9112 bp) using blunt end cloning (BEC) and Gibson assembly, respectively. The resulting sequences can be found as pU3U5_miniHIV_4p8kbp_seq.txt, pU3U5_restrictedto3p4kbp_seq.txt, and pU3U5_enlargedto9p1kbp_seq.txt as part of the zip archive Plasmid_data_seq.zip in the Supporting Information. All three plasmids exhibit two 180 bp terminal sequences corresponding to viral U3 and U5 ends. The plasmids are linearized using the cutting enzyme ScaI. AFM samples are prepared by incubating 5 ng/μl DNA and IN at different concentrations in high sodium buffer (HSB; 250 mM NaOAc, 25 mM Tris, 5 mM Mg(OAc)_2_; pH=7.6) at room temperature, and subsequent 5-fold dilution with 25 mM Tris. The sample was deposited on freshly cleaved mica modified with aminopropylsilatrane (APS)(*26*). The sample was incubated for 30 s before being washed with 20 ml of MilliQ water and dried with a gentle stream of filtered argon gas.

After sample preparation, AFM images were acquired in tapping mode at room temperature using a Nanowizard Ultraspeed 2 (JPK, Berlin, Germany) AFM with silicon tips (FASTSCAN-A, drive frequency 1400 kHz, tip radius 5 nm, Bruker, Billerica, Massachusetts, USA). Images were scanned over (5 × 5) μm^2^ or smaller areas at a line scan speed of 5 Hz. The free amplitude was adjusted to the complex heights, usually 20 nm. The amplitude set point was set to ∼80% of the free amplitude and adjusted to maintain good image resolution.

Force-volume experiments were performed in aqueous buffer (50 mM NaOAc, 25 mM Tris, 1 mM Mg(OAc)_2_; pH= 7.6) using the Nanowizard Ultraspeed 2 (JPK, Berlin, Germany) this time with silicon tips (BL-AC40TS, drive frequency 25 kHz in water, tip radius 10 nm, Olympus, Tokyo, Japan). Cantilever spring constants and optical lever sensitivities were determined by fitting the frequency spectrum of thermal fluctuations, using a Lorentzian function. Imaging uses a peak force of 300 pN, loading rate of 2 μm/s, and a sampling rate of 20 kHz. Images were scanned over different fields of view and with pixel sizes between 4 and 10 nm (indicated for each image).

### AFM data analysis

The post-processing of the AFM data was carried out using SPIP software (v.6.4, Image Metrology, Hørsholm, Denmark) to smooth and line-correct the images. Next, the images were analyzed using custom-written Python code (available from the authors upon request) to determine the radii of gyration for the core of the IN-DNA clusters and the clusters themselves.

### Analysis of AFM based force-volume data

Processing of force volume maps was performed using MountainsSPIP v9 (Digital Surf) software and includes baseline z-correction using a first order polynomial, and x-alignment of the curves using the approach segment of the force curves. The contact point for x-alignment is taken the first crossing of the deflection (y-) axis. The measured piezo height is converted to lateral tip position by correction for the bending of the cantilever. To this end the cantilever deflection (measured in units of length) is subtracted from the piezo height.

Individual force curves were analyzed in terms of stiffness, elastic deformation energy, and plastic deformation energy. Stiffness *k* was determined by taking the slope of a linear fit to the approach segment within the force regime of 0-200 pN. Elastic deformation energy *E_Elast._*, (defined as the area enclosed by the retract segment above the x-axis, i.e. at zero force), and the plastic deformation energy *E_Plast._* (defined as the area enclosed by the approach segment above the y-axis), were used to define a viscoelasticity index 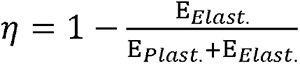.

Maps of the spatially resolved stiffness and deformation energies were generated and used to identify and extract areas corresponding to nucleoprotein complexes. The pixel values were pooled per particle and the midpoint of the Abott-Firestone curve (i.e. at 50% material ratio) was used to determine means and standard deviations. Significance tests used two-sample t-test.

### Magnetic tweezers setup

Force spectroscopy experiments were performed on a custom-built MT setup described previously(*51*). Two magnets (5 × 5 × 5 mm^3^; W-05-N50-G, Supermagnets, Switzerland) are mounted vertically(*34*, *52*) on a motorized arm with a translational motor (M-126.PD2 motor with C-863.11-Mercury controller, PI, Germany) and a rotational motor (C-150.PD motor with C-863.11-Mercury controller, PI, Germany) to control the rotation and z-position of the magnets. The outlet of the flow cell is connected via an adapter to a peristaltic pump (ISM832C, Ismatec) for fluid flushing. The setup is controlled by Lab-VIEW software (National Instruments) described previously (*53*). The flow cells were constructed from two glass coverslips (24 × 60 mm, Carl Roth, Germany). The lower coverslip was pre-functionalized with (3-glycidoxypropyl)trimethoxysilane (abcr GmbH, Germany) as described(*54*). The silanized slides were then covered with 75 μL of a 5000-fold diluted stock solution of polystyrene beads (Polysciences, USA) in ethanol (Carl Roth, Germany) and slowly air dried to later serve as reference beads for drift correction. The upper coverslip was equipped with two holes, ∼1 mm in diameter, drilled with a laser cutter to allow for fluid exchange in the flow cell. The lower and upper coverslips were assembled with a single layer of parafilm (Carl Roth, Germany), pre-cut to form a ∼50 μL channel connecting the inlet and outlet ports of the flow cell, on a heating plate at 80 °C for 30s. After flow cell preparation, 100 µl of 200 µg/mL anti-digoxigenin (Abcam, Germany) in 1× phosphate-buffered saline (PBS) was introduced into the flow cell and incubated overnight. The flow cell was then rinsed with 800 µL of 1× PBS and passivated with BSA (25 mg/mL; Carl Roth, Germany) for 1 hour to minimize non-specific interactions. The flow cell was then rinsed with 800 µL of 1× PBS to remove any residual material.

### DNA constructs and magnetic beads for magnetic tweezers

For the MT experiments, we used a 21-kb DNA construct, prepared as described previously (*55*). The two ends of the DNA segments were provided with handles (∼600 bp) containing several biotins and several digoxigenins, respectively, to bind magnetic beads and the bottom surface of the flow cell. The magnetic beads used were streptavidin-coated MyOne beads of 1 μm diameter (Thermo Fisher Scientific, Germany). The DNA construct was attached to the beads by incubating 0.5 µL picomolar DNA stock solution and 2 µL of MyOne beads in 200 µL PBS for 30 s. The DNA bead solution was then introduced into the flow cell to allow the multiple digoxigenin-anti digoxigenin bonds to form on the surface. The flow cell was then rinsed with 1 mL PBS to remove any unbound beads.

### Magnetic tweezers measurements protocol

Before introducing IN to the flow cell, DNA-bound beads were examined for the presence of multiple DNA tethers by measuring their response to force and torque. To find out whether a magnetic bead is bound to the surface by more than one DNA tether, we introduce negative rotations under high tension (F = 5 pN). In the case of a single double-stranded DNA, this level of force prevents the formation of plectonemes upon underwinding due to DNA melting, and consequently no height change is observed. In contrast, if a bead is bound to two or more double-stranded DNA molecules, braiding occurs when the bead is rotated, resulting in a decrease in z-extension. Beads bound by multiple tethers are excluded from further analysis.

The length of the DNA tethers was checked by varying the force from 5 pN to <0.1pN. At 5 pN the DNA constructs are stretched to >95% of the contour length, whereas at <0.1 pN they drop to the surface and the extension is close to zero. Only the tethers of a length close to the expected contour length (∼7 µm) were considered for the experiment. Once the DNA tethers for the experiment were selected, the flow cell was flushed with 800 µl of low sodium buffer (LSB, 25 mM Tris-HCl pH= 7.6; 5 mM MgCl_2_; 50 mM NaCl; pH= 7.6), followed by 50 µl of 2 mM IN in LSB flushed at 1.7 µl/min, while holding the magnet to exert a force of 5 pN onto the beads. Subsequently, the force was lowered to have a control value of the tether length at 1pN for the analysis. The magnet was next moved to exert a very low force (>0.1 pN) in order to allow for integrase-DNA interaction. Force-jump measurements then involved magnet movement after 5 min of incubation, to cause a sudden force jump to 1 pN. The z-extension of the DNA tethers was monitored over 3-4 h or overnight. For force-extension measurements, the magnet is moved stepwise to induce force-increments of 0.1 pN (for forces smaller than 1 pN) and 0.5 pN (for forces larger than 1 pN). Extension-time traces are recorded at each force plateau for 60 min (for forces smaller than 1 pN) and 30 min (for forces larger than 1 pN).

### Analysis of magnetic tweezers data

Magnetic tweezers data were analyzed using custom written code in Matlab (R2022a, Mathworks). Force-extension measurements were analyzed by correcting the time traces for drift by subtracting the (*x,y,z*) time traces of surface attached reference beads. For each force plateau, the mean extension was computed and plotted against previously calibrated forces. To enable averaging over multiple beads and account for different offsets, we normalize the extension with respect to the end-end distance for each bead at *F* = 5 pN. In order to identify deviations from the worm-like chain induced by IN, we compare the data in presence of IN to the fitted (*56*) force extension behavior in the absence of IN.

For force-jump measurements, the individual tethers time traces were drift corrected using the *x,y,z*-coordinates of reference beads, de-spiked by linear interpolation using the *filloutliers* function in Matlab, and smoothed using a Butterworth filter. The resulting extension-time traces are numerically differentiated to yield velocity-time traces, and subsequently analyzed using the peak finder function. Peaks in the velocity data correspond to “extension bursts” where the effective DNA length increases, due to de-compaction of the DNA from DNA-IN clusters. Dwell times between bursts are calculated from successive peak locations, burst velocities are evaluated at the peak maximum, and burst sizes are calculated by peak integration.

### Monte Carlo simulations of DNA-IN interactions and conformations

We simulated the DNA-IN systems using coarse-grained Monte Carlo (MC) simulations. We refer to reference (*57*) for further details. In the simulations, we modelled the DNA with a bead size of 12 bp of DNA. The DNA beads interact via an excluded-volume interaction to prevent overlapping and subsequent beads are connected by springs. To reproduce the experimentally determined bending stiffness of DNA, we introduce a bending potential, which penalizes sharp bends between subsequent springs. The IN proteins are modelled as beads of the same size as the DNA beads. The IN beads correspond to tetramers and can bind to DNA beads via an isotropic, short-ranged binding potential, where the tetrameric nature of IN is captured by imposing a maximum valence of 4, i.e. each IN bead can bind to at most four DNA beads. To determine the binding strength between IN and DNA, we systematically simulated the binding probability of IN to DNA as a function of the binding strength. These data are then compared to experimental data, where we have directly determined the binding probability from AFM images (Supporting Figure Fig. S22). Briefly, we quantified the fraction of IN-DNA complexes with respect to the total number of binding sites (assuming a site size of 16 bp, the linear DNA has 1000 – 16 + 1 = 985 sites per molecule). We quantify the fraction of bound sites in the presence of 100 nM and 200 nM IN. The best fitting binding energy of IN to DNA was determined to be 5 *k*_B_*T* (Fig. 3 in the main text).

In order to investigate the role of the IN-IN interaction in the IN-induced compaction of DNA, we considered two cases for IN-IN interaction: (i) a purely excluded-volume interaction (same as for the DNA beads), and (ii) an excluded-volume interaction with an added short-ranged attraction. For both cases, we performed multiple simulations for a range of IN concentrations and DNA lengths. In all simulations, the DNA was simulated in the canonical ensemble, while we simulated the IN in the grand-canonical ensemble to ensure that the free IN is kept constant at the value assigned at the start of the simulation. From the simulations, we obtained time series showing different compaction pathways. To classify these pathways and the individual states along these pathways, we developed an unsupervised machine learning pipeline. For this we composed a training dataset of roughly 15,000 snapshots from the simulation time series. Then, for each snapshot in this dataset, we computed 15 different parameters which captured the structural configuration of the DNA-IN complex, including the radius of gyration of the DNA chain, the DNA bend angles, the number of proteins bound to the DNA, and the number and size of protein clusters. Next, we used principal component analysis (PCA) to reduce the dimensionality of the 15-dimensional input. We found that 3 dimensions were sufficient to retain all important features of the input data. Furthermore, we observed distinct clusters in this 3-dimensional PC distribution. Therefore, we used a clustering algorithm –specifically a Gaussian mixture model (GMM)– to provide an unsupervised and objective clustering of the 3-dimensional space into seven categories for (compacted) DNA-IN complexes. Using the trained unsupervised machine learning pipeline, we then classified every configuration of each time series.

### Data analysis and statistics

AFM force-volume data we statistically assessed using a 2-sample *t*-test. Statistical significance of the effect of inhibitor CX014442 was tested using 2-sample Kolmogorov-Smirnov test. All statistical tests were performed in Matlab (Mathworks). *P* values of <0.05 were considered significant.

## Acknowledgments

We thank Thomas Nicolaus for laboratory assistance, and Enrico Skoruppa, Enrico Carlon, Michael Hafner, Frank Smallenburg, Chris Brackley, Rinske Alkemade, Gerhard Blab, Wolfgang Ott, Paulo Onuchic, and Arthur Ermatov for helpful discussions.

## Author contributions

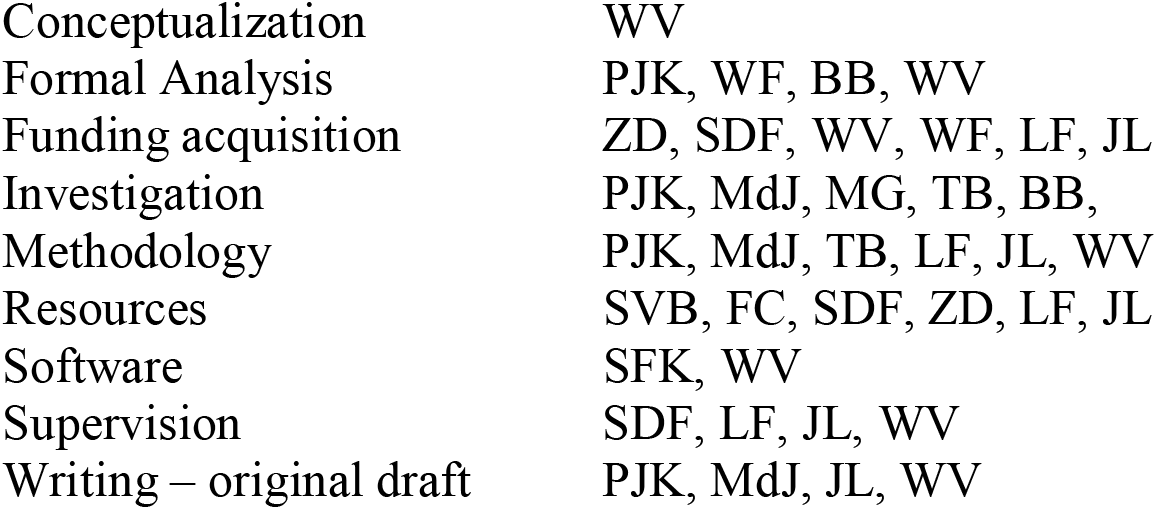

## Data and materials availability

All data are available in the main text or the supplementary materials.

## Supplementary Materials

Supporting Figures S1-S22.

Supporting Data S1

