## Supplemental material for "HIV integrase compacts viral DNA into biphasic condensates"

Pauline J. Kolbeck *et al.*

**This PDF file includes:**

Figs. S1 to S22

Data S1

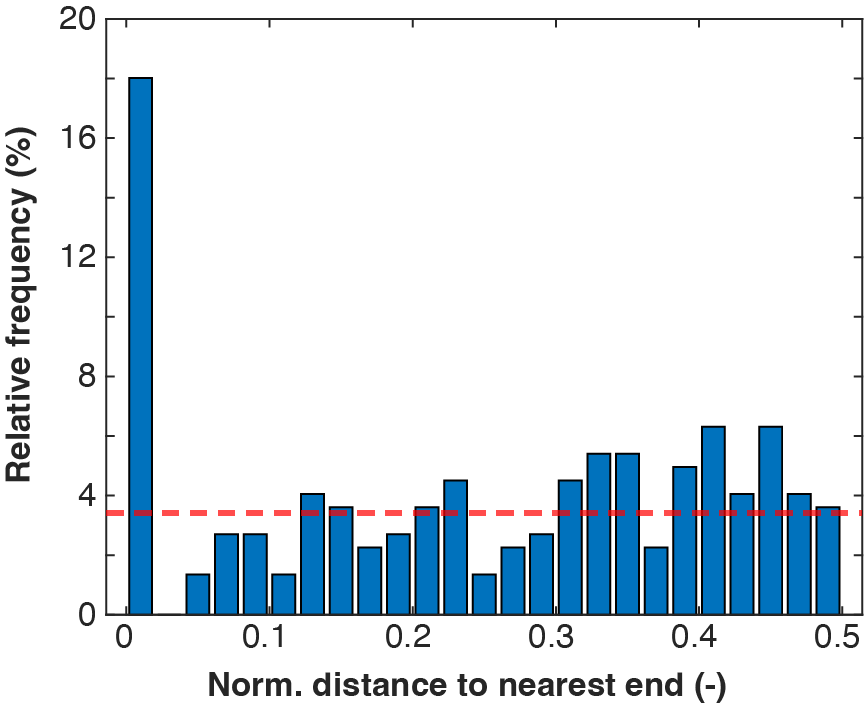

Fig. S1.

**Distribution of binding positions of integrase on a 491 bp DNA construct.** For the measurement, blunt end DNA in Mg^2+^-buffer was used. The red line indicates the average relative frequency for completely non-specific binding.

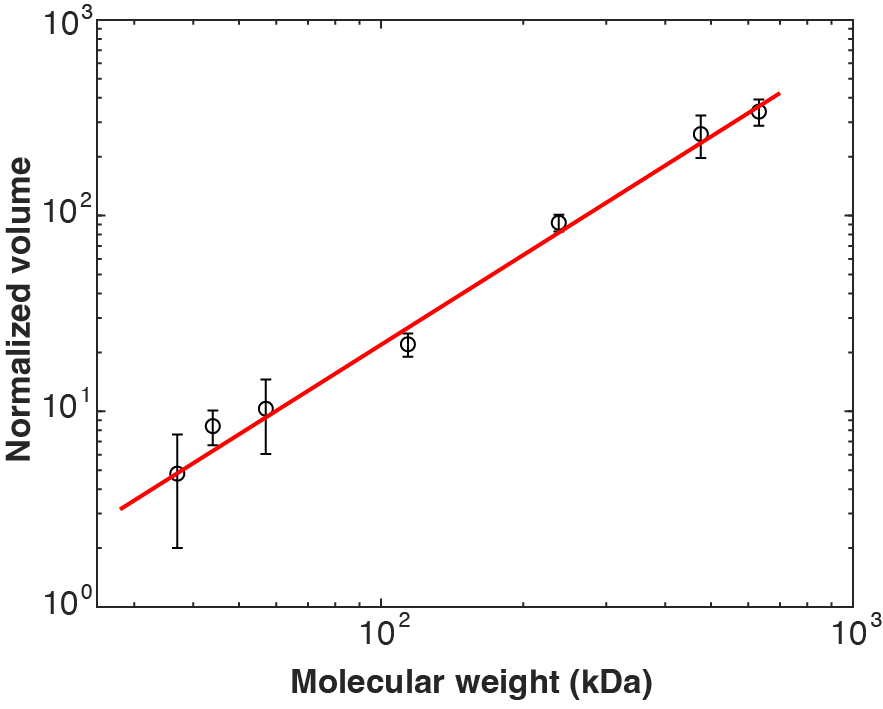

Fig. S2. AFM volume analysis of proteins. Calibration curve of normalized volumes (mean ± SD) of PvuII (dimer 36kDa), Prototype foamy virus integrase (monomer 44.4 kDa), EcoRV (dimer 57.2 kDa, tetramer 114.4 kDa), UvrA-mCherry fusion (dimer 238 kDa, tetramer 476 kDa) and maltose binding protein-β-galactosidase fusion (tetramer 632 kDa).

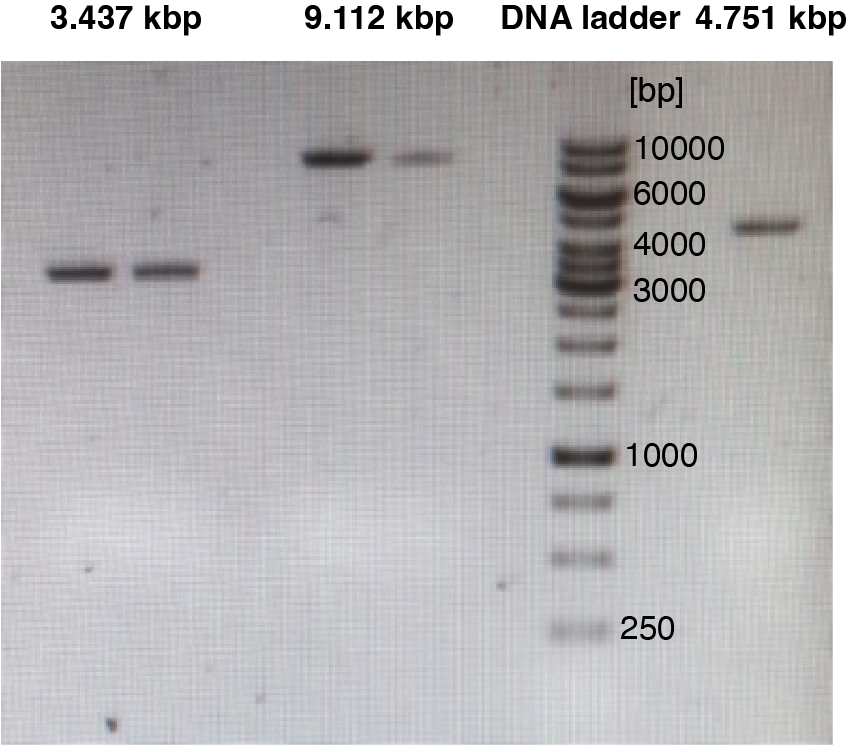

Fig. S3. Gel electrophoresis of the different length DNA constructs used in this study. The lanes are from left to right: 3437 bp DNA (two different Mini-Preps); empty lane; 9112 bp DNA (two different Mini-Preps); empty lane; DNA ladder (1 kbp gene ruler, Thermo Fisher); empty lane; 4751 bp DNA.

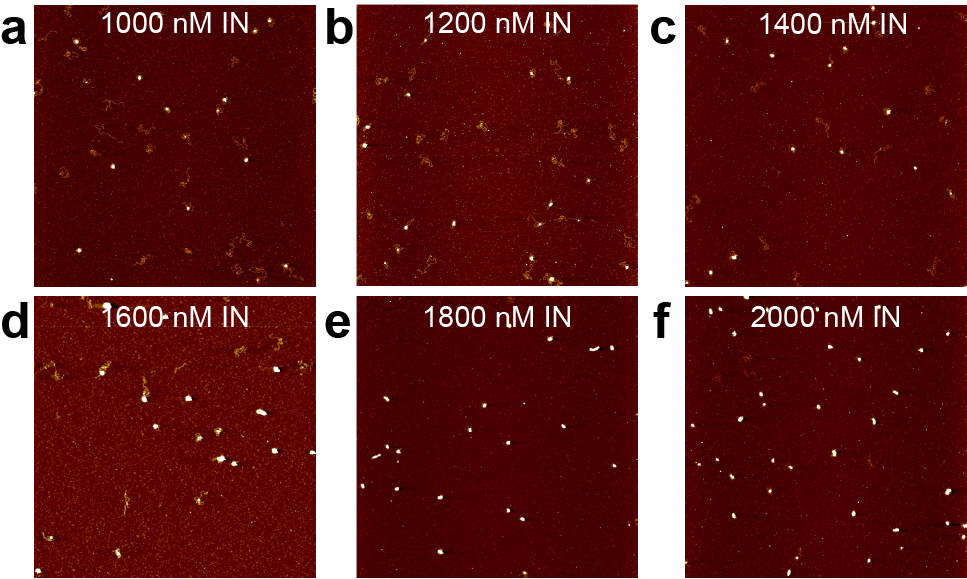

**Fig. S4**. **AFM images of 3.4 kbp DNA compaction by IN.** The compaction of 3.4 kbp DNA is very similar to 4.8 kbp DNA except for the fact that there is no rosette state observed and the compaction occurs in one single step. a. – f. Overview AFM topographs (5 μm x 5 μm) of the 3.4 kbp DNA in the presence of IN in concentrations ranging from 1000 nM to 2000 nM.

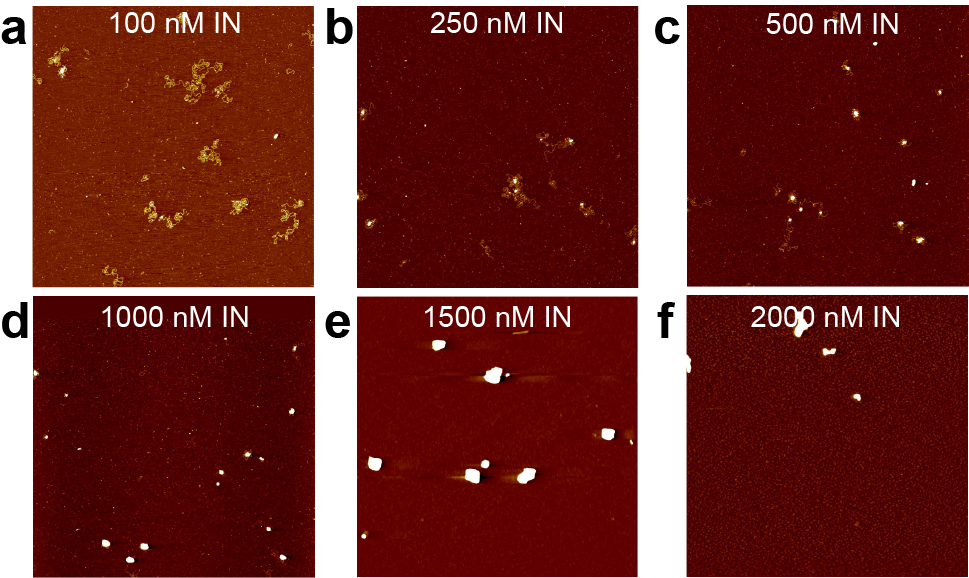

**Fig. S5. AFM images of 9.1 kbp DNA compaction by IN.** The compaction of 9.1 kbp DNA is very similar to 4.8 kbp DNA. **a. – f.** Overview AFM topographs (5 μm x 5 μm) of the 9.1 kbp DNA in the presence of IN in concentrations ranging from 100 nM to 2000 nM.

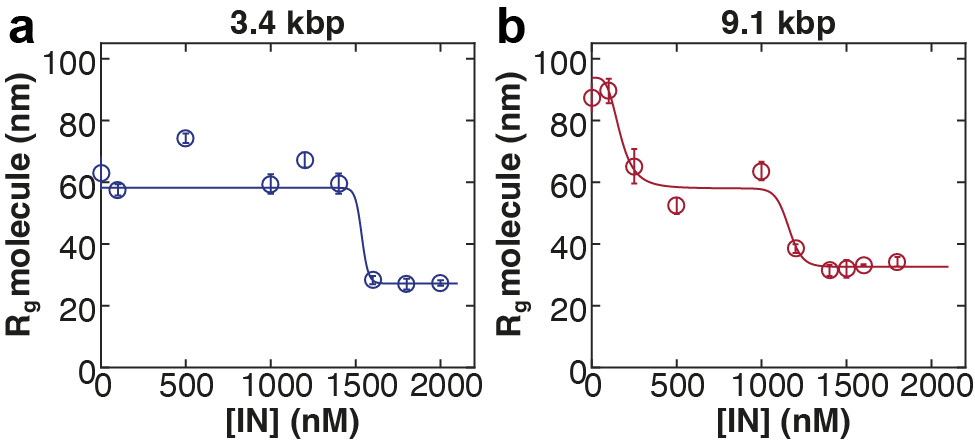

**Fig. S6. Radii of gyration of the nucleoprotein complex for 3.4 and 9.1 kbp DNA at 60 mM ionic strength as a function of [IN].** **a.** The data for the 3.4 kbp DNA (see Supplementary Fig. S4) reveal only one compaction transition at ~ 1500 nM. It is therefore fitted by a single-Hill-fit. **b.** The data for the 9.1 kbp DNA (see Supplementary Fig. S5) show two distinct compaction transitions and are fitted by a double-Hill-fit: the first transition from open state to rosette state occurs at ~ 300 nM, the second transition from rosette state to fully compacted state at ~ 1100 nM.

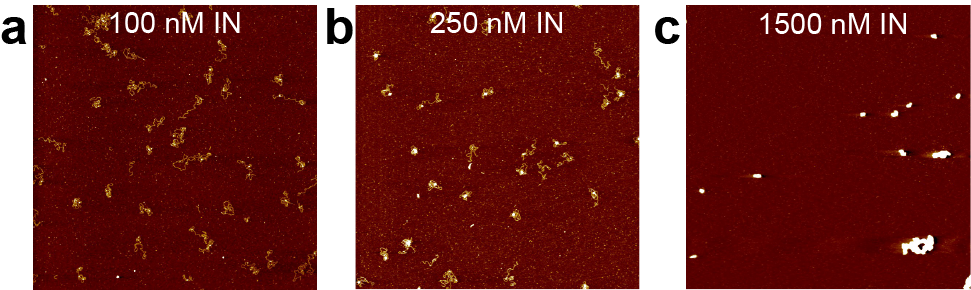

**Fig. S7. AFM images of DNA incubated with IN at 37 °C instead of at room temperature. a. – c.** Overview AFM topographs (5 μm x 5 μm) of the 4.8 kbp DNA in the presence of 100, 250, and 1500 nM IN incubated at 37 °C for 30 minutes before deposition. We find very similar complexes as are observed at the same concentration after incubation at room temperature. Since no effect of the incubation temperature on the compaction behavior was observed, the incubation was carried out at room temperature for all experiments shown in this work for convenience.

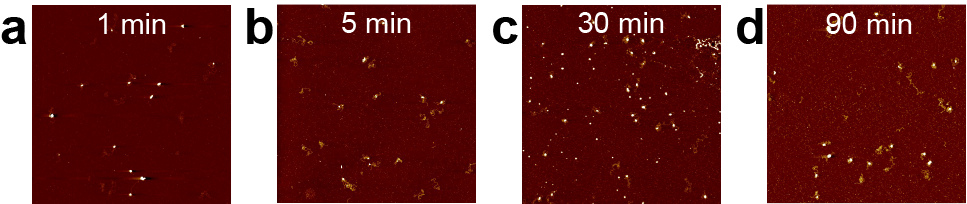

**Fig. S8. AFM images of DNA and IN after different incubation times. a. – d.** Overview AFM topographs (5 μm x 5 μm) of the 4.8 kbp DNA in the presence of 500 nM IN for different incubation times (1 min – 90 min) before deposition. We do not observe systematic difference with incubation time over the range of times probed. Since no effect of the incubation time on the compaction behavior was observed, the incubation time was set to 5 min for all experiments shown in this work for convenience.

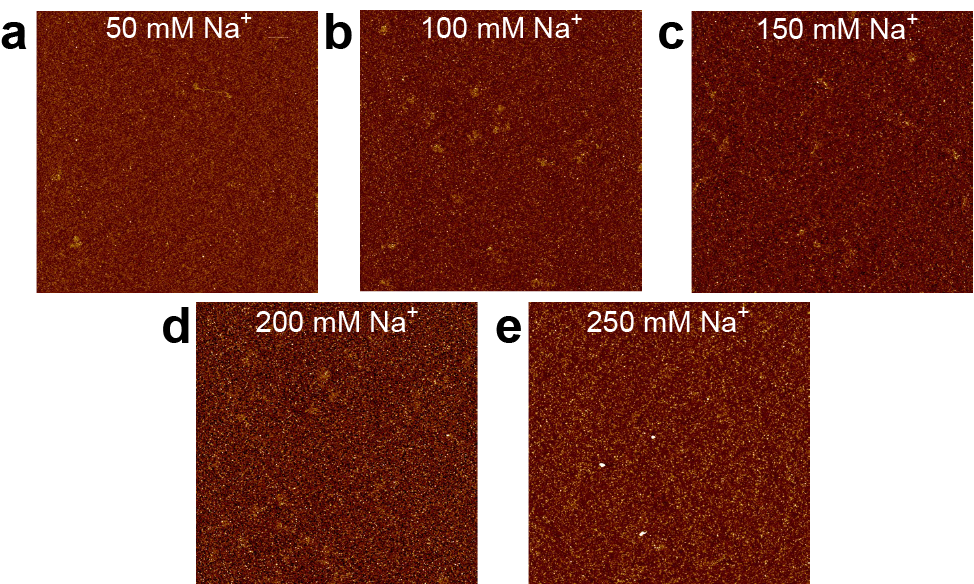

**Fig. S9. AFM images of DNA incubated with 2000 nM IN without a dilution step at different sodium concentrations. a. – e.** Overview AFM topographs (5 μm x 5 μm) of the 4.8 kbp DNA in the presence of 2000 nM IN at sodium concentrations ranging from 50 mM to 250 mM. Without the incubation step, no DNA compaction is observed.

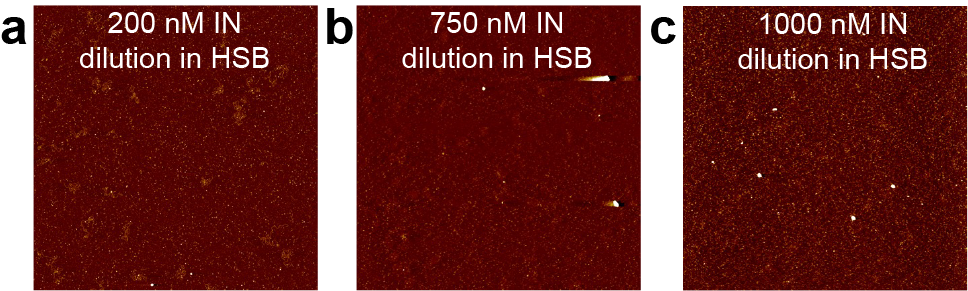

**Fig. S10. AFM images of DNA incubated with IN with a dilution step but at different ionic strengths. a. – c.** Overview AFM topographs (5 μm x 5 μm) of the 4.8 kbp DNA in the presence of varying IN concentrations (200, 750, and 1000 nM). After incubation, the DNA-IN mix is not diluted with Tris or LSB (50 mM Na^+^, 5mM Mg^2+^) as done for the other experiments, but instead with HSB (250 mM Na^+^, 5mM Mg^2+^). No DNA compaction is observed.

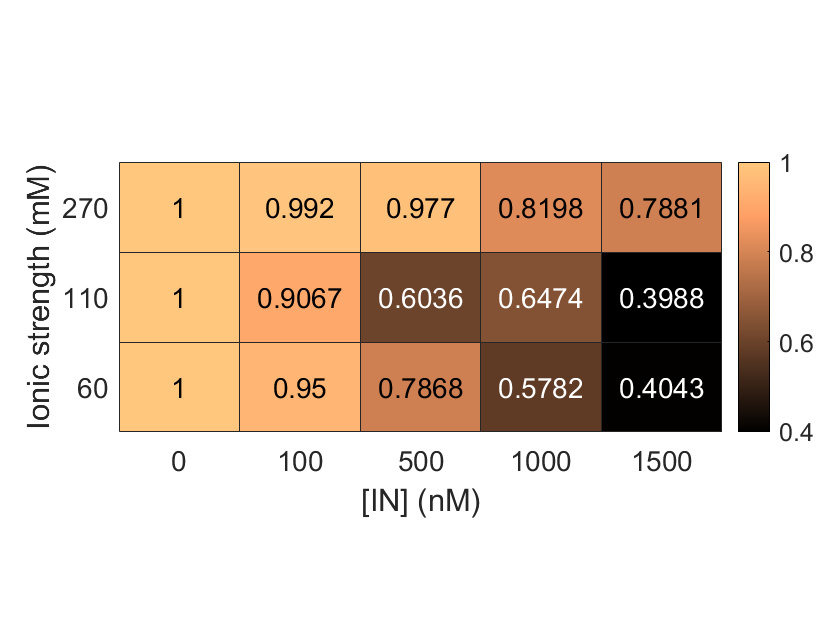

**Fig. S11. Phase diagram depicting the normalized radii of gyration of the nucleoprotein complexes formed as a function of [IN] and ionic strength of the solution.** At high ionic strength, compaction is significantly less than at lower ionic strength and no full compaction is reached even at high [IN].

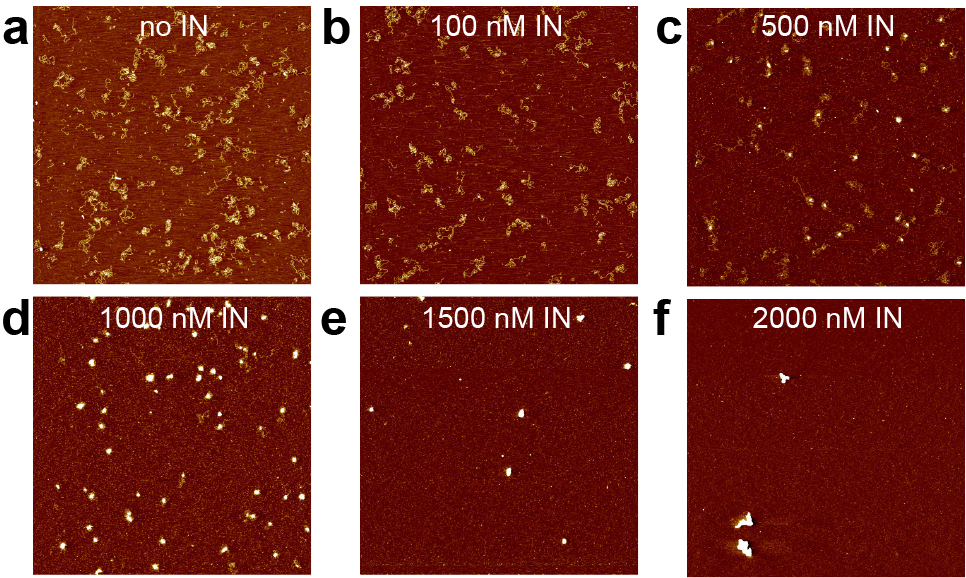

**Fig. S12. AFM images of 4.8 kbp DNA with unspecific ends at different IN concentrations. a. – f.** Overview AFM topographs (5 μm x 5 μm) of a 4.8 kbp DNA constructs with unspecific ends (i.e. not using the viral cDNA sequences at the ends) in the presence of IN in concentrations ranging from 1000 nM to 2000 nM. The compaction of 4.8 kbp DNA with unspecific ends is very similar to 4.8 kbp DNA with 180 bp ends identical to the viral DNA ends.

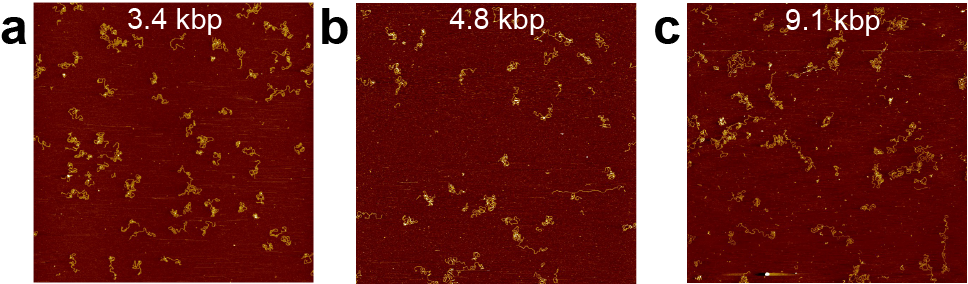

**Fig. S13. AFM images of bare DNA. a.** Overview AFM topograph (5 μm x 5 μm) of the 3.4 kbp DNA without IN. **b.** Same as in panel a for 4.8 kbp DNA. **c.** Same as in panel a for 9.1 kbp DNA. For all three DNA lengths, no compaction is observed in the absence of IN but otherwise identical conditions (buffer contains 50 mM Na^+^, 1 mM Mg^2+^).

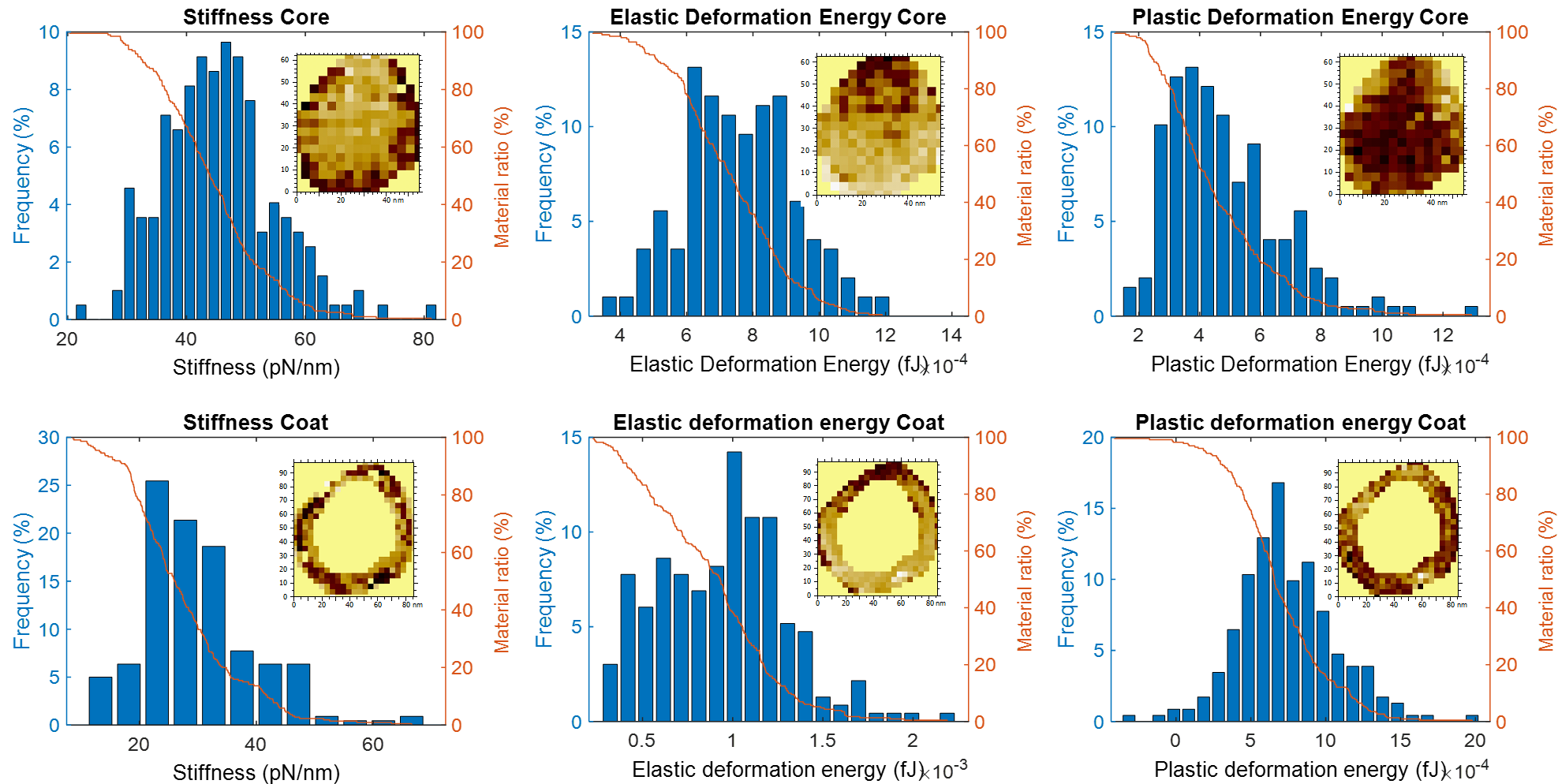

**Fig. S14. Multiparametric imaging of DNA-IN complexes and quantification of elastic and viscoelastic constants.** Distributions (blue) and Abott-Firestone curves (orange) of the values for indentation stiffness, elastic deformation energy, and plastic deformation energy (left to right) for the core and coat (top and bottom panels, respectively) of a fully compacted complex. Insets show the corresponding parameter maps. The data points in Fig. 4e,f were determined from the 50% points in the Abott-Firestone plots.

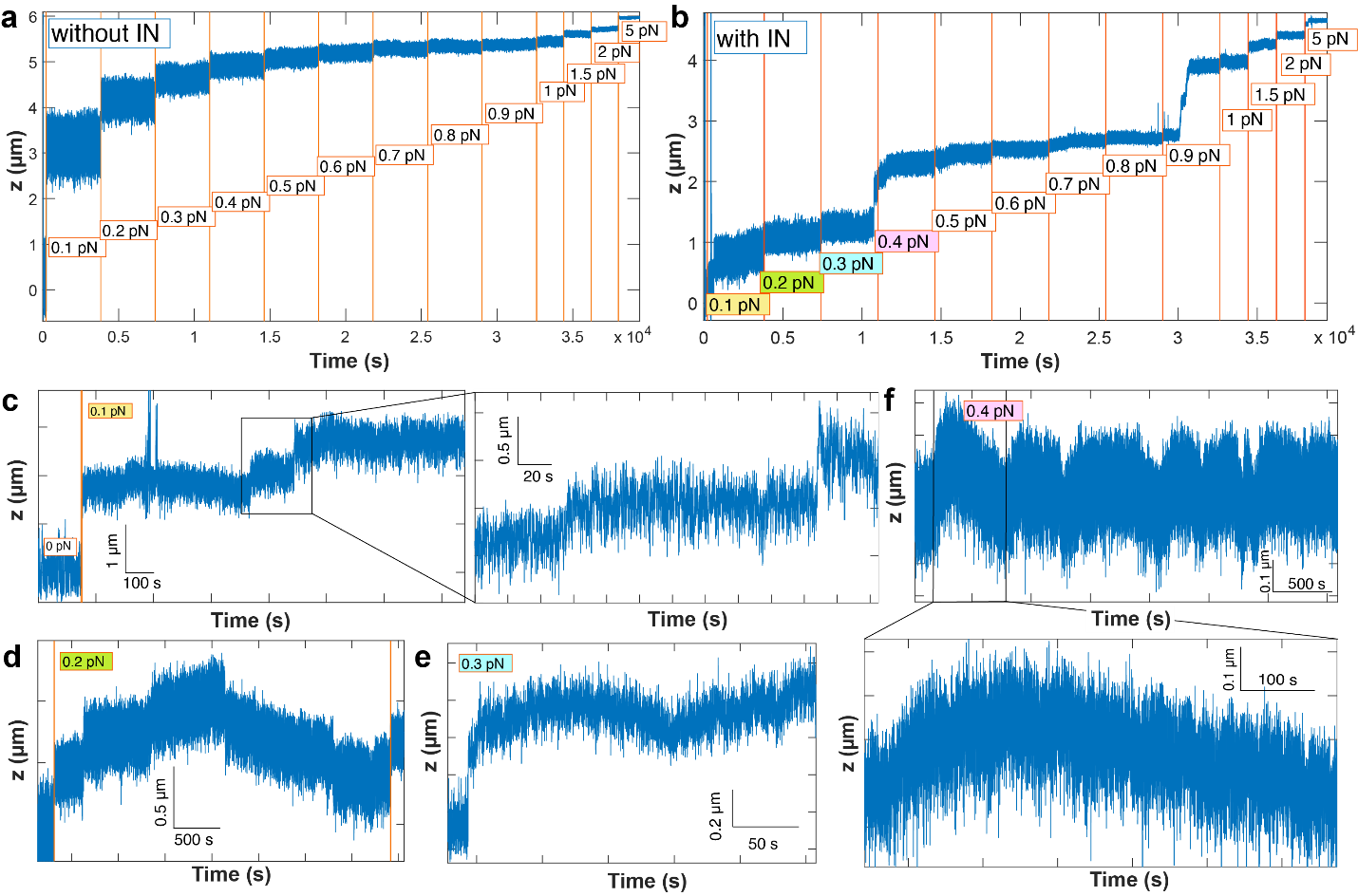

**Fig. S15. Exemplary magnetic tweezers (MT) extension time traces at different forces. a.** MT extension time traces at forces ranging from 0.1 to 5 pN recorded for bare DNA in the absence of IN. **b.** MT extension time traces at forces ranging from 0.1 to 5 pN recorded in the presence of 2 μM IN. **c.** Extension trace at 0.1 pN in the presence of 2 μM IN and an additional zoom to visualize the broad fluctuations. **d.** Extension trace at 0.2 pN in the presence of 2 μM IN, revealing changes in length corresponding to de-compaction and compaction of several hundred nanometers. **e.** Extension trace at 0.3 pN in the presence of 2 μM IN. **f.** Extension trace at 0.4 pN in the presence of 2 μM IN and an additional zoom to visualize the continuous compaction and de-compaction at constant force. The traces for the individual force plateaus in panel c-f are not from the same data set shown in panel b.

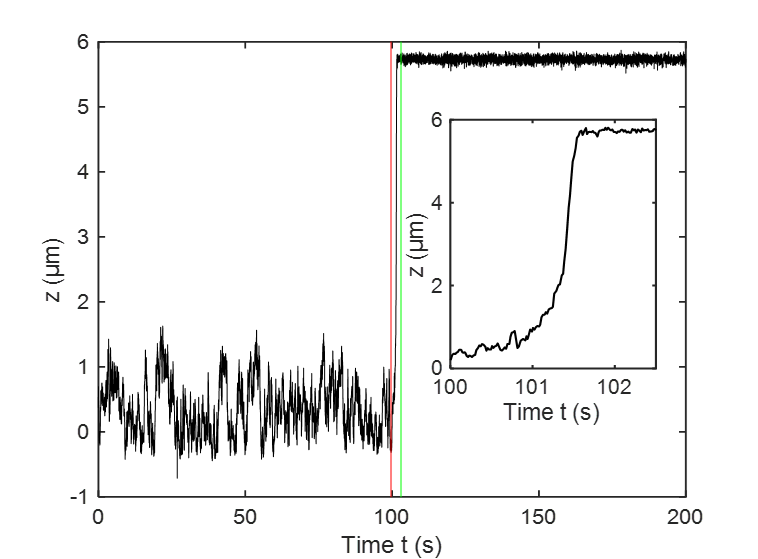

**Fig. S16. Magnetic tweezers force-jump control experiment in the absence of IN.** Extension-time trace for a force jump from 10 fN to 1 pN (at *t* ≈ 100 s) in the absence of IN. The trace shows a very rapid extension increase upon increasing the force within the time of external magnet movement (<1 s).

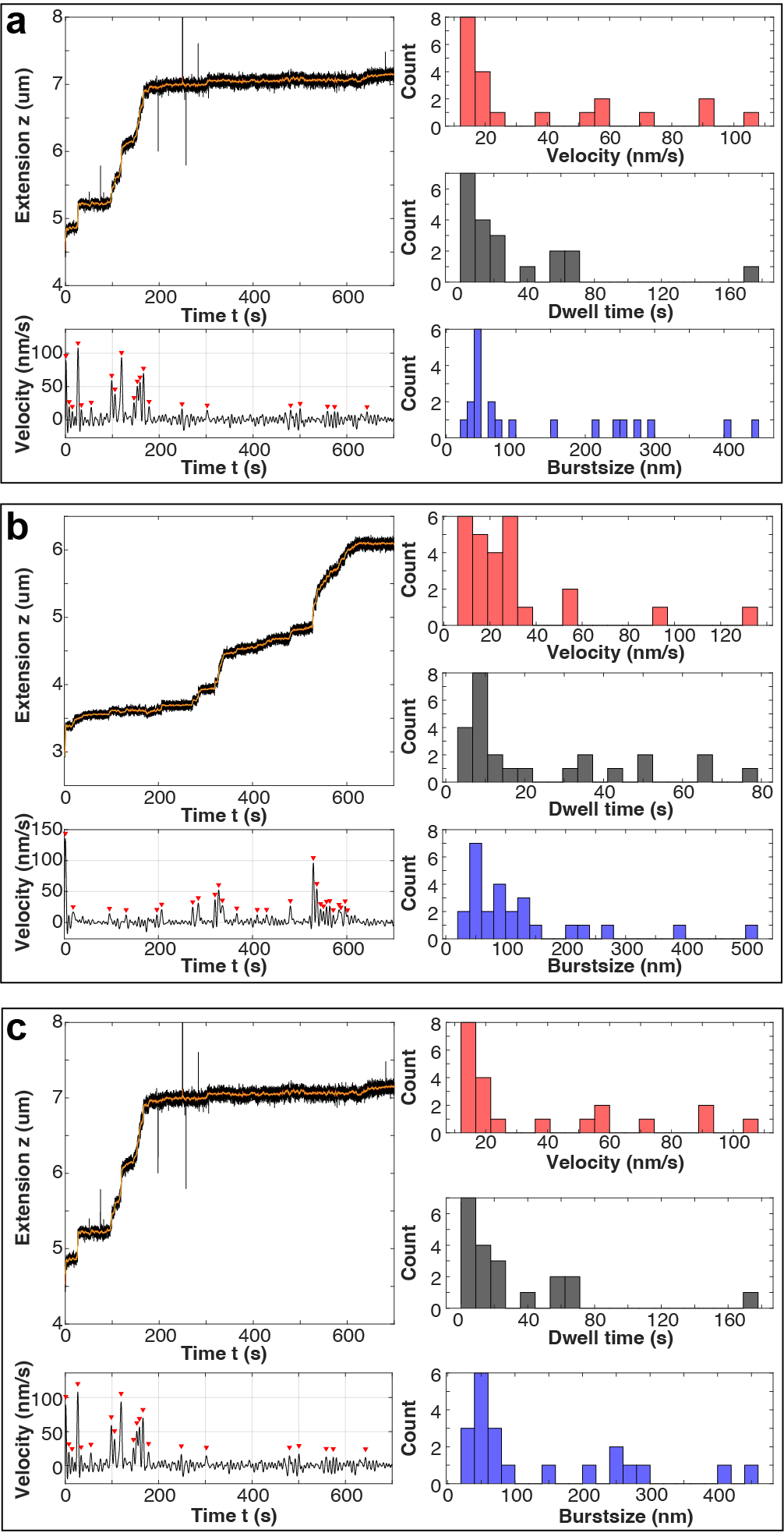

**Fig. S17. Examples of MT extension time traces and their analysis. a,b.** Two exemplary extension time traces at a force of 1 pN (top left of each panel). Black traces are the raw data recorded at 72 Hz. Red traces are processed data to remove spikes (due to tracking errors) and to reduce fluctuations (due to thermal motion of the bead-tether system) using a Butterworth filter (see Methods). Further analysis of the filtered extension-time traces includes subsequent numerical differentiation to yield velocity-time traces and velocity peak detection (bottom left). Peaks are analyzed in terms of their amplitude (i.e. their local maximum; top right), the time between peaks (i.e. dwell times; middle right), and the area under the velocity peaks (i.e. the extension increment or burst size; bottom right).

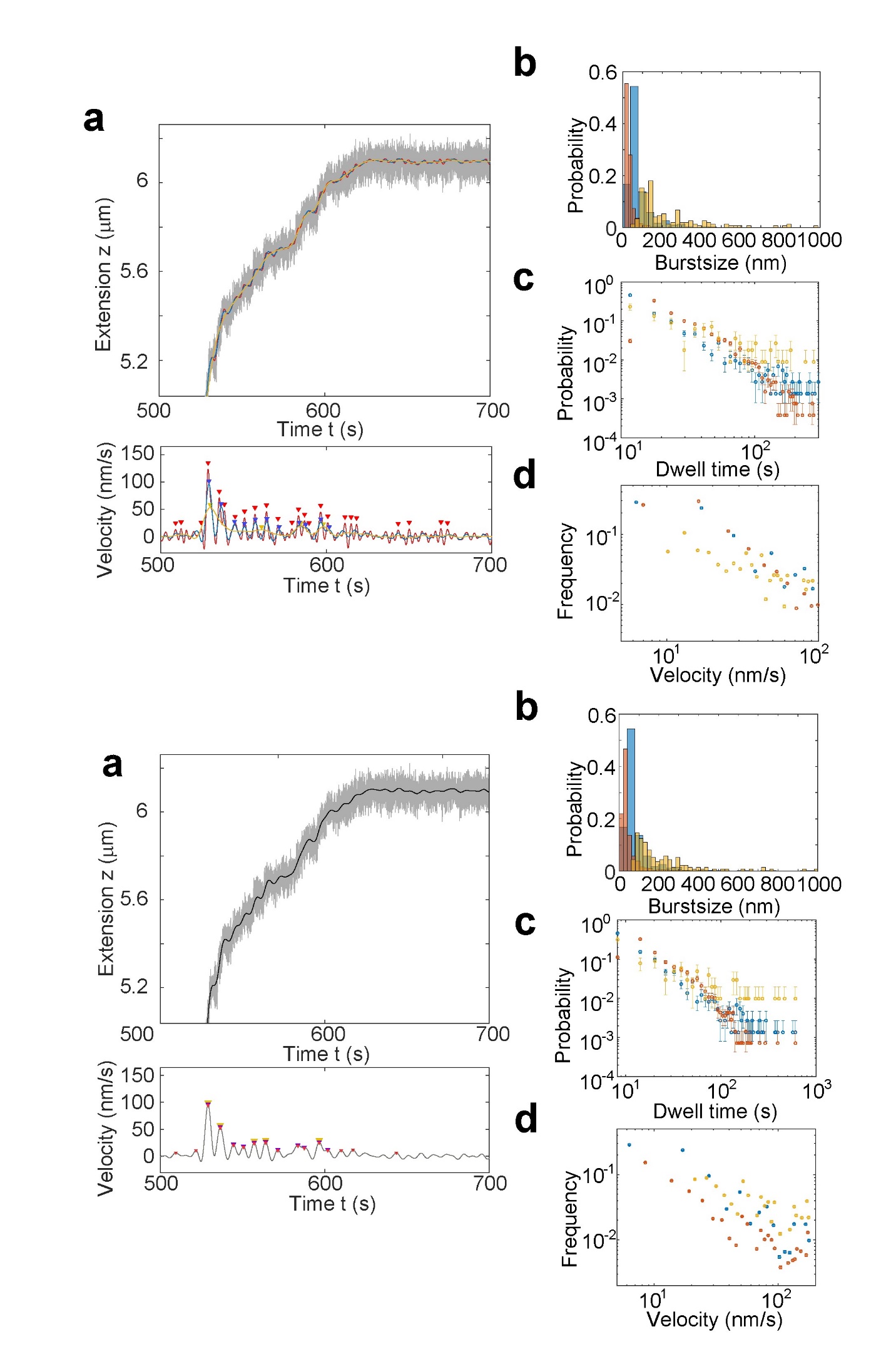

**Fig. S18. Effect of the threshold for burst detection at constant filter settings for extension time traces. a.** Top plot: Extension versus time trace showing raw data (72 Hz, grey) and smoothed data (Butterworth filter with a half power frequency 0.0025; black). Bottom plot: velocity time trace obtained by numerical differentiation of the extension trace, with detected velocity peaks at cutoff value 5.75 nm/s (red triangles) 11.5 nm/s (blue triangles), and 23 nm/s (yellow triangles). **b.** Burst size distributions for cutoff values of 5.75 nm/s (red) 11.5 nm/s (blue), and 23 nm/s (yellow). **c.** Dwell time distributions for cutoff values of 5.75 nm/s (red) 11.5 nm/s (blue), and 23 nm/s (yellow). **d.** Weighted velocity distributions for cutoff values of 5.75 nm/s (red) 11.5 nm/s (blue), and 23 nm/s (yellow).

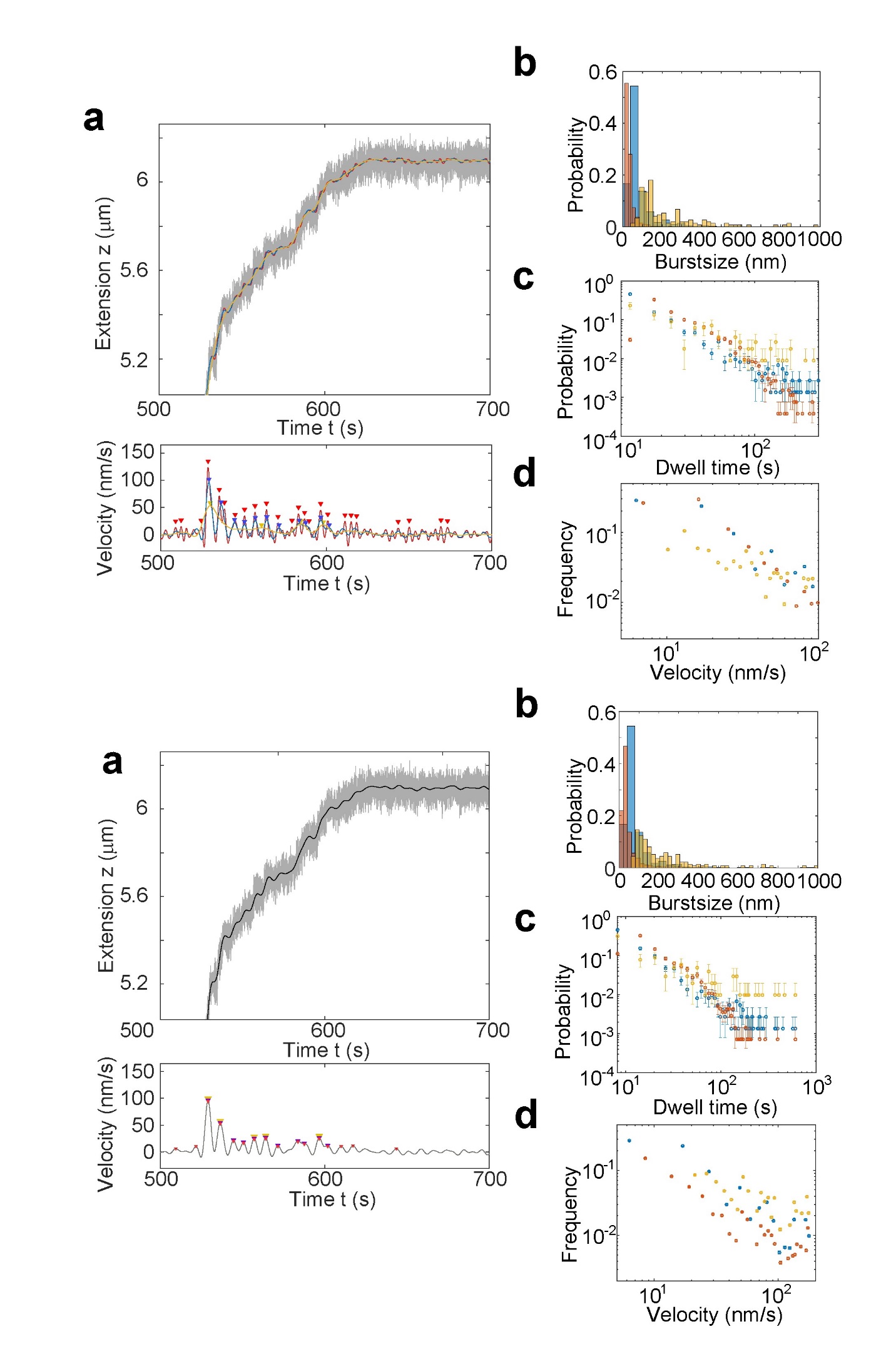

**Fig. S19. Effect of filter settings for burst detection at constant threshold value for extension time traces. a.** Top plot: Extension versus time trace showing raw data (72 Hz, grey) and smoothed data (Butterworth filter with half power frequency 0.0025 (red), 0.005 (blue), and 0.01 (yellow). Bottom plot: velocity-time trace showing detected velocity peaks at cutoff value 11.5 nm/s for the different half power frequencies applied **b.** Burst size distributions for half power frequency of 0.0025 (red), 0.005 (blue), and 0.01 (yellow). **c.** Dwell time distributions for half power frequencies of 0.0025 (red), 0.005 (blue), and 0.01 (yellow). **d.** Weighted velocity distributions for half power frequencies of 0.0025 (red), 0.005 (blue), and 0.01 (yellow).

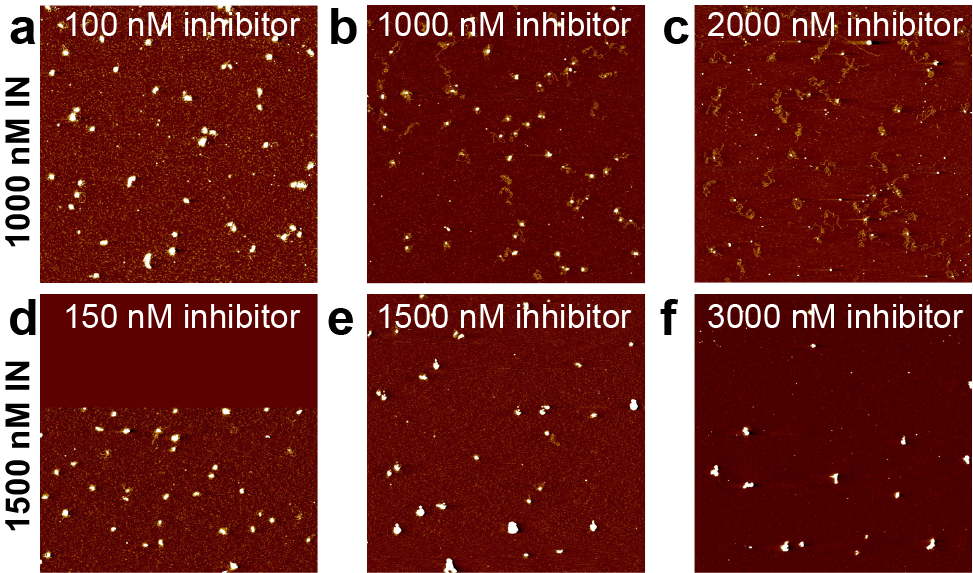

**Fig. S20. AFM images probing the effect of adding inhibitors on 4.8 kbp DNA compaction by IN. a. – f.** Overview AFM topographs (5 μm x 5 μm) of the 4.8 kbp DNA in the presence of 1000 nM (top row) or 15000 nM IN (bottom row), respectively, and increasing inhibitor concentration from left to right (as indicated in the individual panels). The inhibitor was added after complex formation. The compaction of 4.8 kbp DNA is significantly reduced by the addition of increasing concentrations of inhibitor for the rosette state (1000 nM IN), but largely unaffected for the fully compacted state (1500 nM IN; see also Fig. 6 in the main text).

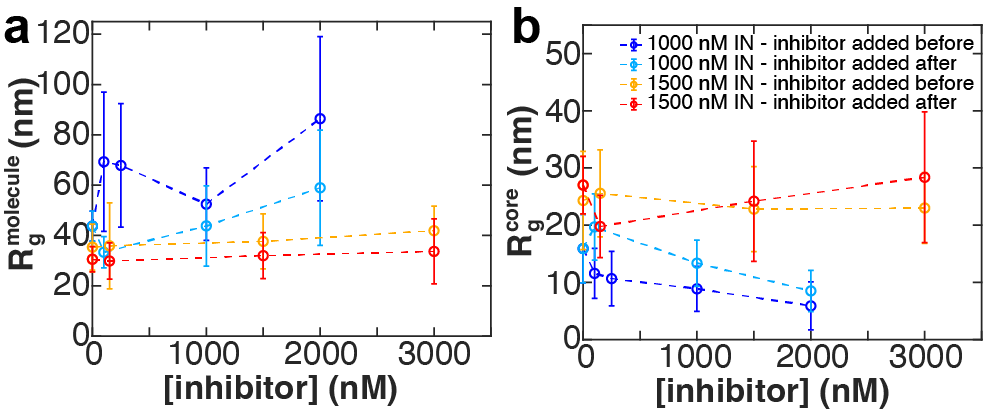

**Fig. S21. Effect of adding inhibitors to IN-DNA condensates before and after compaction. a.** *R_g_*-values for the entire molecule for the rosette state (1000 nM IN; dark and light blue) and the fully compacted state (1500 nM IN; orange and red) for varying [inhibitor] and for adding the inhibitors before or after the compaction reaction. Symbols are the mean and standard deviations from at least 20 molecules. **b.** Same as in panel a for the *R_g_*-values of the core of the condensates.

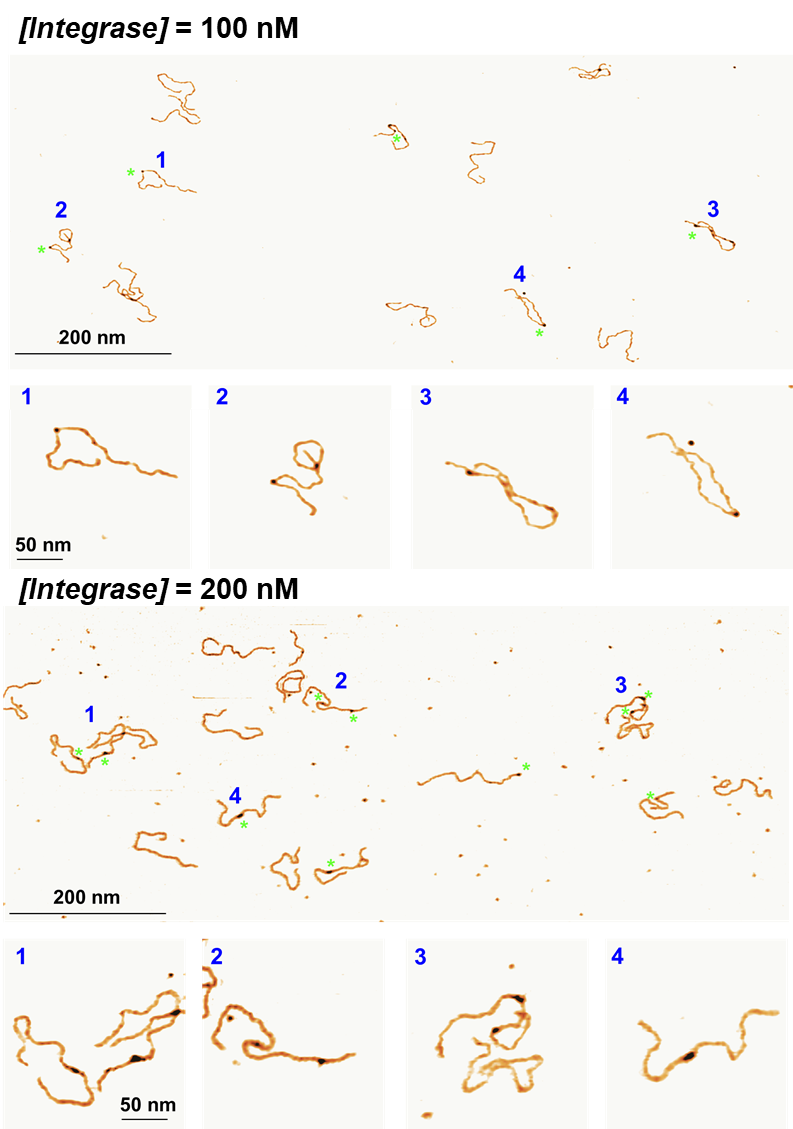

**Fig. S22. Experimental data used for determining IN-DNA affinity.** AFM images of 1000 bp HIV DNA mimetic in the presence of 100 nM and 200 nM integrase final concentration. We quantify the fraction of DNA binding sites bound over the total number of sites, assuming a site size of 16 bp. Our quantification is based on 910 DNA molecules in the presence of 100 nM IN, and 1796 DNA molecules in the presence of 200 nM IN.

Data S1. (separate file)

Sequences of plasmids used or created in this work are provided as .txt files in zip file “Plasmid_data_seq”: pU3U5_miniHIV_4p8kbp_seq.txt; pU3U5_miniHIV_enlarged to 9p1kbp_seq.txt; pU3U5_miniHIV_restrictedto3p4kbp_seq.txt
